# *Alu* insertion-mediated dsRNA structure formation with pre-existing *Alu* elements as a novel disease-causing mechanism

**DOI:** 10.1101/2020.01.30.926790

**Authors:** Emmanuelle Masson, Sandrine Maestri, Valérie Bordeau, David N. Cooper, Claude Férec, Jian-Min Chen

## Abstract

We previously identified a homozygous *Alu* insertion variant (*Alu*_Ins) in the 3’-UTR of the *SPINK1* gene as the cause of a novel pediatric disease. Although we established that *Alu*_Ins leads to the complete loss of *SPINK1* mRNA expression, the precise mechanisms remained elusive. Here we aimed to elucidate these mechanisms through a hypothesis-driven approach. Initially, we speculated that *Alu*_Ins could independently disrupt mRNA 3’ end formation and/or affect other post-transcriptional processes such as nuclear export and translation, due to its particular location. However, the presence of *Alu*_Ins in the 3’-UTR resulted in only an ~50% reduction in luciferase reporter activity compared to the wild-type, suggesting the involvement of additional mechanisms. Using RepeatMasker, we identified two *Alu* elements within *SPINK1*’s third intron, both of which resided in an orientation opposite to that of *Alu*_Ins. Through RNAfold predictions and full-length gene expression assays designed to examine orientation-dependent interactions between *Alu* repeats, we present evidence linking the detrimental effect of *Alu*_Ins to extensive double-stranded RNA structures formed between *Alu*_Ins and pre-existing intronic *Alu* sequences. Our results reveal a novel pathogenetic mechanism involving an *Alu* insertion, highlighting the importance of considering interactions between new and pre-existing *Alu* elements in inverted orientations within disease-associated genes.

## 1. Introduction

*Alu* elements are short DNA sequences approximately 300 nucleotides in length, representing one of the most abundant types of mobile genetic element within primate genomes. They exceed one million copies in the human genome, constituting nearly 11% of the total genomic content [1]. *Alu* elements are continually amplified in the human genome through long interspersed element-1 (LINE-1 or L1)-mediated retrotransposition [2, 3], a mechanism also linked to various human genetic disorders (for reviews, see [2, 4-6]). Historically, disease-causing *Alu* insertions have invariably been identified within the coding or proximal intronic regions of affected genes, largely due to detection bias [7]. However, a new finding emerged in 2017 when a full-length *Alu* insertion, henceforth termed *SPINK1 Alu*_Ins, was identified for the first time outside of these conventional gene regions — it was found in the 3’-untranslated region (3’-UTR) of the *SPINK1* gene (MIM: 167790) in a patient presenting with severe infantile isolated exocrine pancreatic insufficiency, a novel pediatric disease [8]. This novel finding would not have been possible without the inclusion of the gene’s 3’-UTR in the mutational screen. Additionally, the *Alu*_Ins variant might easily have been overlooked had it not been present in the homozygous state. More recent examples of *Alu* insertions identified outside the conventional coding or proximal intronic regions of disease genes include an *Alu*Jb element within the *SMN1/2* promoter region [9] and a 316 bp *Alu* insertion overlapping the *TMEM106B* 3’-UTR that is tightly linked with top GWAS variants associated with the increased risk of frontotemporal lobar dementia with TDP-43 inclusions (FTLD-TDP) [10].

The pathological implications of disease-causing *Alu* insertions within coding or proximal intronic regions are typically linked to their potential to disrupt coding sequences or to induce aberrant splicing, often leading to a lack of detailed experimental analysis. In our previous research, we explored the functional consequences of the *SPINK1 Alu*_Ins variant using a cell culture-based full-length gene expression assay (FLGEA) [8] (see also Figure 1). We found that, whereas the wild-type (WT) *SPINK1* expression vector produced normally spliced transcripts, the mutant (Mut) vector resulted in a complete absence of *SPINK1* mRNA expression in transfected HEK293T cells. This finding was supported by three distinct lines of evidence: First, reverse transcription-polymerase chain reaction (RT-PCR) analysis detected *SPINK1* transcripts in lymphocytes from a healthy control but failed to identify any *SPINK1* transcripts in lymphocytes from the *SPINK1 Alu*_Ins homozygote [8]. Second, a complete deletion of the entire *SPINK1* gene in the homozygous state, leading to complete loss of *SPINK1* expression, was found in another patient with severe infantile isolated exocrine pancreatic insufficiency [8]. Third, there were notable pathological similarities between the pancreatic conditions in these two patients and those in *Spink3*-deficient mice [11] (Note: *Spink3* is the murine equivalent of *SPINK1*). Importantly, the pancreatic conditions resulting from complete *SPINK1* functional loss were significantly more severe than those in *SPINK1* c.194+2T>C homozygotes with chronic pancreatitis, who retained about 10% of normal *SPINK1* function [12-14].

**Figure 1.**
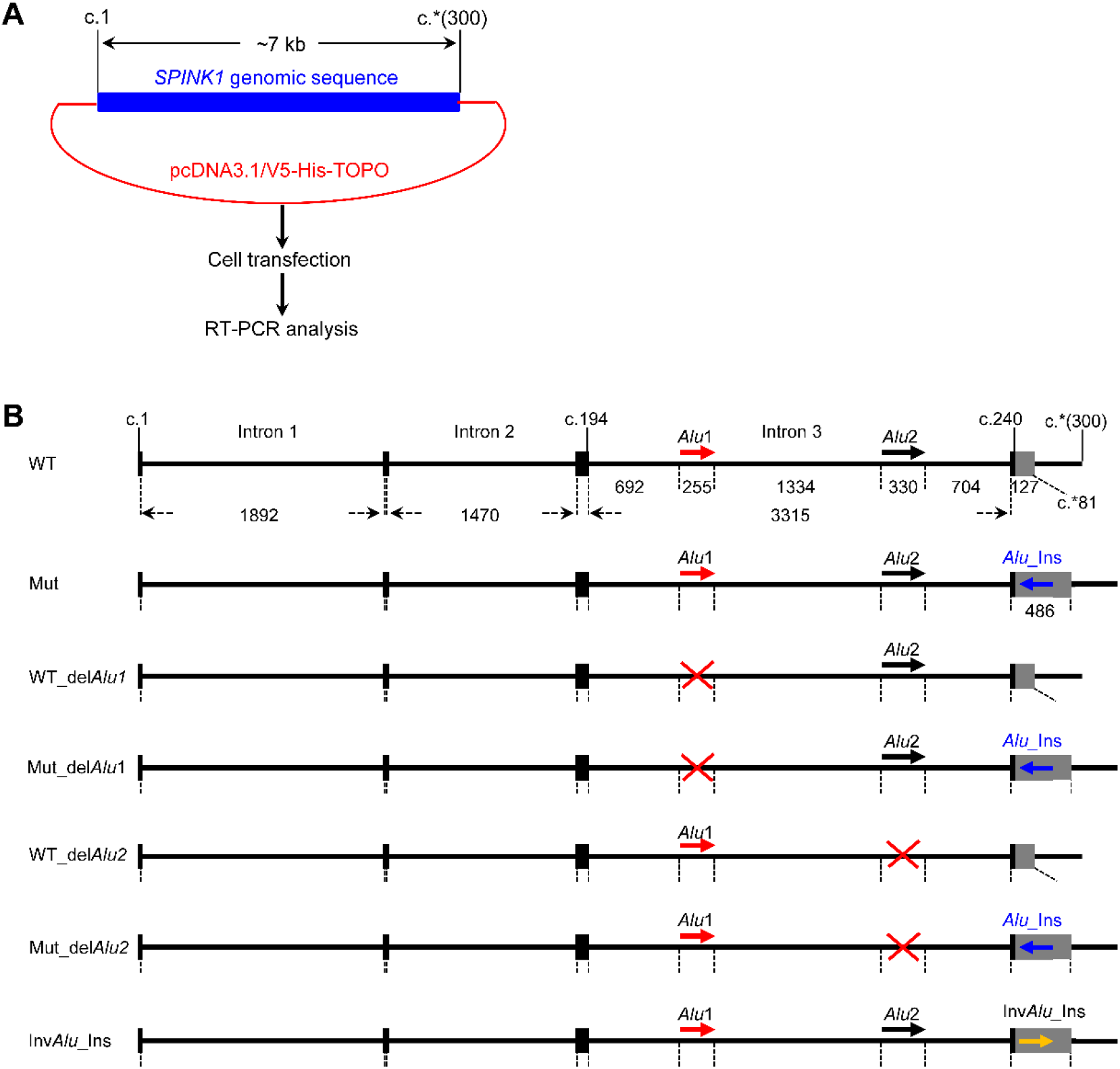
Illustration of the FLGEA assay and the seven expression vectors used in the current study. (**A**) Schematic representation of the expression vectors and the experimental procedures of the FLGEA assay. Each expression vector was constructed using the pcDNA3.1/V5-His TOPO vector as a backbone. The *SPINK1* sequences inserted in these vectors span from c.1 (the A of the ATG start codon) to c.*(300) (300 bases beyond the TGA stop codon), based on the WT *SPINK1* genomic sequence. For further sequence information, see Supplementary Figure S1. (**B**) Detailed depiction of the WT and Mut *SPINK1* sequences and the five novel artificial sequences developed for the FLGEA vectors in this study. The *SPINK1* genomic sequence is shown to scale, with coding regions indicated by black bars and the 3’-UTR as a grey box. The lengths (in bp) of the three introns and the two *Alu* elements (named *Alu*1 and *Alu*2) in intron 3 are specified. Positions c.194, c.240, and c.*81 correspond to the final nucleotide positions of exon 3, the translation termination codon, and the 3’-UTR, respectively. Horizontal solid arrows show the orientations of the pre-existing *Alu* elements in intron 3, the disease-associated *Alu* insertion (*Alu*_Ins), and the artificially inverted version of *Alu*_Ins (Inv*Alu*_Ins), with rightward arrows denoting alignment with the *SPINK1* gene’s sense strand and leftward arrows indicating the opposite orientation. Artificial deletions of *Alu*1 or *Alu*2 in WT and Mut contexts are highlighted with red crosses. *Abbreviations:* FLGEA, full-length gene expression assay; Mut, mutant; RT-PCR, reverse transcription-polymerase chain reaction; WT, wild-type; 3’-UTR, 3’-untranslated region.

Despite these findings, the precise mechanisms underlying the complete loss of *SPINK1* mRNA expression due to *SPINK1 Alu*_Ins have remained elusive. In the present study, we aimed to elucidate these mechanisms through a hypothesis-driven approach.

## 2. Materials and Methods

### 2.1. Reference sequences

In line with our earlier studies [8, 13, 15], the pathophysiologically relevant, four-exon *SPINK1* gene [16, 17] was employed as the reference. The corresponding mRNA reference sequence is NM_001379610.1. The genomic sequence for *SPINK1* was sourced from the GRCh38/hg38 assembly, accessible at https://genome.ucsc.edu/. Nucleotide numbering followed the coding DNA sequence guidelines established by the Human Genome Variation Society [18].

### 2.2. 3’-UTR luciferase reporter assay

#### 2.2.1. Construction of WT and Alu_Ins 3’-UTR reporter vectors

The WT *SPINK1* gene’s 3’-UTR spans 81 bp. To construct the WT 3’-UTR reporter vector, we amplified a 306-bp fragment that includes the entire 3’-UTR (from c.*1 to c.*81) and the downstream 225-bp 3’ flanking sequence (from c.*82 to c.*306) of the *SPINK1* gene (see Supplementary Figure S1). This fragment was PCR-amplified from genomic DNA obtained from a healthy individual using the forward primer 5’-<u>TCTAGA</u>GAACCAAGGTTTTGAAATCCCA-3’ (containing an *Xba*I restriction site, underlined) and the reverse primer 5’-<u>GGATCC</u>GATCATCTGTGCTCTGCCAT-3’ (containing a *BamH*I restriction site, underlined). The corresponding fragment containing the *Alu*_Ins was amplified from genomic DNA obtained from the homozygous patient. The resulting PCR products were firstly digested with *Xba*I and *BamH*I and then cloned into the *XbaI*/*BamH*I sites of the pGL3 Control Vector (Promega, Charbonnieres, France), yielding the pGL3-WT and pGL3-*Alu SPINK1* 3’-UTR reporter gene constructs, respectively. In each construct, the insert was placed immediately after the translational termination codon of the luciferase reporter gene. The accuracy of both inserts was confirmed through sequencing using the BigDye™ Terminator v1.1 Cycle Sequencing Kit (ThermoFisher Scientific, Illkirch, France).

#### 2.2.2. Cell culture, transfection and luciferase reporter gene assay

Human embryonic kidney (HEK293T) and human pancreatic adenocarcinoma (COLO-357) cell lines were maintained in DMEM nutrient mixture supplemented with 10% fetal calf serum. Transfections were carried out as previously described [19] using 3.8 μg pGL3-WT or pGL3-*Alu SPINK1* 3’-UTR luciferase reporter vector plus 0.2 μg control pRL-CMV vector. At 48 h after transfection, luciferase activity was measured as previously described [20].

### 2.3. Identification of pre-existing *Alu* elements in the *SPINK1* locus using RepeatMasker

The search for *Alu* elements in the *SPINK1* locus was conducted using RepeatMasker (http://www.repeatmasker.org/). Two *Alu* elements, identified in intron 3 of the gene, were designated *Alu*1 and *Alu*2 respectively, as illustrated in Supplementary Figure S1 (see also Figure 1).

### 2.4. Construction of five additional expression vectors for FLGEA

In our earlier research [8], we engineered expression vectors for both the full-length WT and the mutant (Mut) *SPINK1* sequences, the latter containing the *Alu*_Ins variant, for the FLGEA assay (Figure 1). In this study, we expanded our toolkit by developing five additional expression vectors tailored for the FLGEA assay. These new vectors are designated as WT_del*Alu*1, Mut_del*Alu*1, WT_del*Alu*2, Mut_del*Alu*2, and Inv*Alu*_Ins, with ‘Inv*Alu*_Ins’ specifically referring to an inversion of the *Alu*_Ins sequence (Figure 1).

#### 2.4.1. Construction of the WT_delAlu1 and Mut_delAlu1 vectors

To create *Alu*1 deletions within WT and Mut *SPINK1* contexts, we used genomic DNA from a healthy control and the patient homozygous for *SPINK1 Alu*_Ins, respectively. Two specific DNA fragments were amplified (color-coded primer pairs shown in Supplementary Figure S2A). For each fragment, a PCR reaction was performed with 50 ng DNA in a 50 μL reaction mixture containing 2.5 U TaKaRa La Taq™ DNA polymerase, 400 μM TakaRa dNTP Mix, and 0.4 μM each of the corresponding primer pair. The PCR program comprised an initial denaturation at 94°C for 1 min, 35 cycles of denaturation at 94°C for 20 s, annealing at 56°C for 20 s and extension at 72°C for 5 min, and a final extension step at 72°C for 10 min. After *Kpn*I restriction enzyme digestion, the two fragments from each DNA source were ligated, then further amplified with primers P1_F and reverse primer P1_R (Supplementary Figure S2A). The PCR was performed using the GoTaq^®^ Long PCR MasterMix (Promega, Charbonnieres, France) according to the manufacturer’s protocol. The PCR program had an initial denaturation at 94°C for 2 min, 35 cycles of denaturation at 94°C for 20 s, annealing at 58°C for 20 s and extension at 65°C for 6 min, and a final extension step at 72°C for 10 min. The final PCR products, WT_del*Alu*1 and Mut_del*Alu*1, were separately cloned into the pcDNA3.1/V5-His TOPO vector according to the manufacturer’s instructions.

#### 2.4.2. Construction of the WT_delAlu2 and Mut_delAlu2 vectors

These vectors were constructed following the same methodology as applied to WT_del*Alu*1 and Mut_del*Alu*1, but they employed distinct sets of primer pairs, as detailed in Supplementary Figure 2B. Furthermore, the PCR amplification extension times were adjusted according to the lengths of the fragments: 6 min for the longer fragment (amplified using P1_F and *Alu*2_*KpnI*_R primers) and 1 min for the shorter fragment (amplified using *Alu2*_*KpnI*_F and P1_R primers).

#### 2.4.3. Construction of the InvAlu_Ins vector

To develop the Inv*Alu*_Ins vector, intended for reversing the orientation of the *Alu*_Ins sequence within the 3’-UTR of the *SPINK1* gene, we first eliminated a pre-existing *Nco*I restriction site found in the third intron of the Mut vector. This alteration was performed using the QuickChange II XL Site-Directed Mutagenesis Kit with the application of specific primers: forward 5’- TGGCCAACATGGTGAAACCCCGTGGTGGCGGGCGCCTATAATAC-3’ and reverse 5’- GTATTATAGGCGCCCGCCACCACGGGGTTTCACCATGTTGGCCA-3’. After modification, three separate fragments were PCR-amplified from the modified plasmid template. Each PCR reaction used 1 ng of the altered plasmid in a 50 μL reaction mix, which included 2.5U TaKaRa La Taq™ DNA polymerase, 400 μM TaKaRa dNTP Mix, and 0.4 μM of each primer in the corresponding pair (Supplementary Figure 2C). The PCR conditions were set as follows: initial denaturation at 94°C for 1 min, followed by 35 cycles of 20 s at 94°C for denaturation, 20 s at 56°C for annealing, and extension at 72°C for 7 min for fragment A and 1 min for fragments B and C. This was concluded with a final extension at 72°C for 10 min. The three resulting fragments were individually digested with *Kpn*I and/or *Nco*I restriction enzymes and subsequently ligated together. The steps following this ligation were conducted in the same manner as outlined earlier for additional vector constructs.

Primer sequences related to the construction of the five new expression vectors are provided in Supplementary Figure 2D. The exon/intron boundaries and ligation junctions within the resulting vector constructs were verified through sequencing.

### 2.5. FLGEA

#### 2.5.1. Cell culture, transfection and reverse transcription (RT)

HEK293T cells were maintained as described in section 2.2.2. For transfections, 1 μg of the corresponding expression plasmid was used per well in a 6-well plate. Four hours prior to RNA extraction, the cells underwent treatment with 50 μg/ml cycloheximide as previously described [21]. Total RNA was extracted from the cells 24 h post-transfection using TRIzol RNA Isolation Reagents (ThermoFisher Scientific). The purity and concentration of the RNA were assessed based on the optical density (OD) measurements at 260 nm and 280 nm.

For RT, 4 μg RNA was first treated with DNase I (ThermoFisher Scientific, Illkirch, France) to eliminate any contaminating DNA. The RT reaction was prepared in a 20 μL volume, including 1 μg DNase I-treated RNA, 10 U RNAse inhibitor (Promega, Charbonnieres, France), 250 ng Oligo(dT) (Qiagen, Courtaboeuf, France), 4 μl 5× First Strand Buffer, 500 μM dNTPs, 5 mM dithiothreitol, and 200 U SuperScript^®^ II Reverse Transcriptase (ThermoFisher Scientific, Illkirch, France). The reaction mixture was incubated at 42°C for 50 min and then deactivated by heating at 70°C for 15 min. After RT, the cDNA was treated with 2 U RNase H (ThermoFisher Scientific, Illkirch, France) to remove any remaining RNA, at 37°C for 20 min.

#### 2.5.2. RT-PCR

RT-PCR was conducted using the forward primer 5’-GAGTCTATCTGGTAACACTGGAGCT-3’ and reverse primer 5’-CAGTCAGGCCTCGCGGTG-3’, employing the GoTaq^®^ Long PCR MasterMix in accordance with the manufacturer’s instructions. The PCR cycle commenced with an initial denaturation at 94°C for 2 min, then proceeded through 40 cycles of 20 s at 94°C for denaturation, 20 s at 58°C for annealing, and 4 min at 65°C for extension. This was concluded with a final extension phase at 72°C for 5 m. The PCR products were subsequently analyzed by electrophoresis on a 1% agarose gel.

#### 2.5.3. Sequencing of the RT-PCR products

The RT-PCR products derived from the WT, WT_del*Alu*1, WT_del*Alu*2, and Inv*Alu*_Ins expression vectors were purified using the Illustra™ ExoProStar™ (GE Healthcare, Orsay, France). Subsequently, they were directly sequenced utilizing the BigDye™ Terminator v1.1 Cycle Sequencing Kit, employing the forward primer 5’-GAGTCTATCTGGTAACACTGGAGCT-3’ and the reverse primer 5’-CAGTCAGGCCTCGCGGTG-3’.

RT-PCR products from the Mut, Mut_del*Alu*1, and Mut_del*Alu*2 expression vectors were individually cloned into the pcDNA3.1/V5-His TOPO vector, following the manufacturer’s guidelines. Transformation employed XL10-Gold Ultracompetent Cells (Stratagene, La Jolla, CA). The transformed cells were plated on LB agar containing 50 mg/mL ampicillin and incubated at 37ºC overnight. Approximately 30 colonies from each transformation were selected for PCR amplification using the GoTaq^®^ Long PCR MasterMix, in line with the manufacturer’s instructions. The PCR utilized forward primer 5’-GGAGACCCAAGCTGGCTAGT-3’ and reverse primer 5’-AGACCGAGGAGAGGGTTAGG-3’, both situated within the vector sequence. The amplification protocol started with an initial denaturation at 94°C for 2 min, followed by 35 cycles of 20 s at 94°C for denaturation, 20 s at 58°C for annealing, and extension at 65°C for 1 min (for Mut and Mut_del*Alu*2) or 4 min (for Mut_del*Alu*1), culminating in a final extension at 72°C for 5 min. PCR products were assessed via electrophoresis on a 1% agarose gel, then purified using Illustra™ ExoProStar™, and sequenced employing the BigDye™ Terminator v1.1 Cycle Sequencing Kit. Sequencing utilized primers 5’- GGAGACCCAAGCTGGCTAGT-3’ (forward), 5’-TGAAAATCGGTGAGTACA-3’ (forward), 5’- GAAAACATCATGAGCATG-3’ (forward), and 5’-AGACCGAGGAGAGGGTTAGG-3’ (reverse). For the larger band (>2 kb) derived from Mut_del*Alu1*, additional sequencing employed three forward primers: 5’- CTGAGATTGACTTGAT-3’, 5’-TCTGAAACCTCCGAGT-3’, and 5’-CTAACTTAAATGTGGCT-3’.

Bands whose identities remained uncertain following the initial sequencing were excised from the agarose gel and purified using the MinElute Gel Extraction Kit (Qiagen, Courtaboeuf, France). These purified fragments were then sequenced using the BigDye™ Terminator v1.1 Cycle Sequencing Kit. The sequencing employed forward primer 5’-GAGTCTATCTGGTAACACTGGAGCT-3’ and reverse primer 5’- CAGTCAGGCCTCGCGGTG-3.

#### 2.5.4. Quantification of RT-PCR bands using ImageJ

The intensities of the RT-PCR bands of interest were quantified with ImageJ.JS software, accessible at https://ij.imjoy.io/.

#### 2.5.5. Additional RT-PCR analysis

For the purpose of amplifying potential transcripts that retain intron 3 but correctly exclude introns 1 and 2, as detected in the FLGEA assay using the Mut vector, we employed various primer combinations for additional PCR analyses on the above prepared cDNA. The three utilized forward primers were: one located in the promoter region of the pcDNA3.1 vector (5’-TGGTGGAATTGCCCTTATGA-3’), one spanning the junction between *SPINK1* exons 1 and 2 (5’-GAGTCTATCTGGTAACACTGGAGCT-3’), and one spanning the junction between *SPINK1* exons 2 and 3 (5’-TGGGAAGAGAGGCCAAATGT-3’). The three reverse primers were all positioned within intron 3, specifically between the *Alu*1 and *Alu*2 regions (5’- CCAAGAAGAAGGTTTGCATTAC-3’, 5’-CTTGGAAATTTCCCAAGTCTC-3’, and 5’- GCATAAACATAGAACATTTCCATC-3’).

### 2.6. Prediction of RNA secondary structures using RNAfold

RNA secondary structures were predicted using the RNAfold tool, which is part of the Vienna RNA Websuite (http://rna.tbi.univie.ac.at/), under default parameters [22]. The analysis covered sequences from the start of exon 3 to the end of the 3’-UTR (i.e., c.*81 in the context of the WT *SPINK1* gene). For the exact sequences of the five alleles (WT, Mut, Inv*Alu*_Ins, Mut-del*Alu*1, and Mut_del*Alu*2) used in the secondary structure prediction, see Supplementary Figures S3-S7.

## 3. Results

### 3.1. Evaluation of the hypothesis: independent impact of *SPINK1 Alu*_Ins on gene expression

The 3’-UTRs of human genes serve critical roles in regulating mRNA 3’ end formation, stability, degradation, nuclear export, subcellular localization, and translation processes [23-26]. Building on this understanding, we initially hypothesized that the *SPINK1 Alu*_Ins variant might independently influence one or more of these regulatory functions due to its particular insertion location. This could potentially explain the previously observed complete absence of *SPINK1* mRNA expression [8]. To investigate this hypothesis, we conducted a 3’-UTR luciferase reporter assay to assess the overall impact of the *SPINK1 Alu*_Ins variant on the aforementioned regulatory processes. The 3’-UTR vector containing the *Alu*_Ins variant resulted in a significant decrease in luciferase activity, demonstrating approximately a 50% reduction compared to the wild-type control in both HEK293T and COLO-357 cell lines (Figure 2). However, this reduction alone does not fully account for the total absence of *SPINK1* expression observed previously [8], suggesting the presence of additional mechanisms that warrant further exploration.

**Figure 2.**
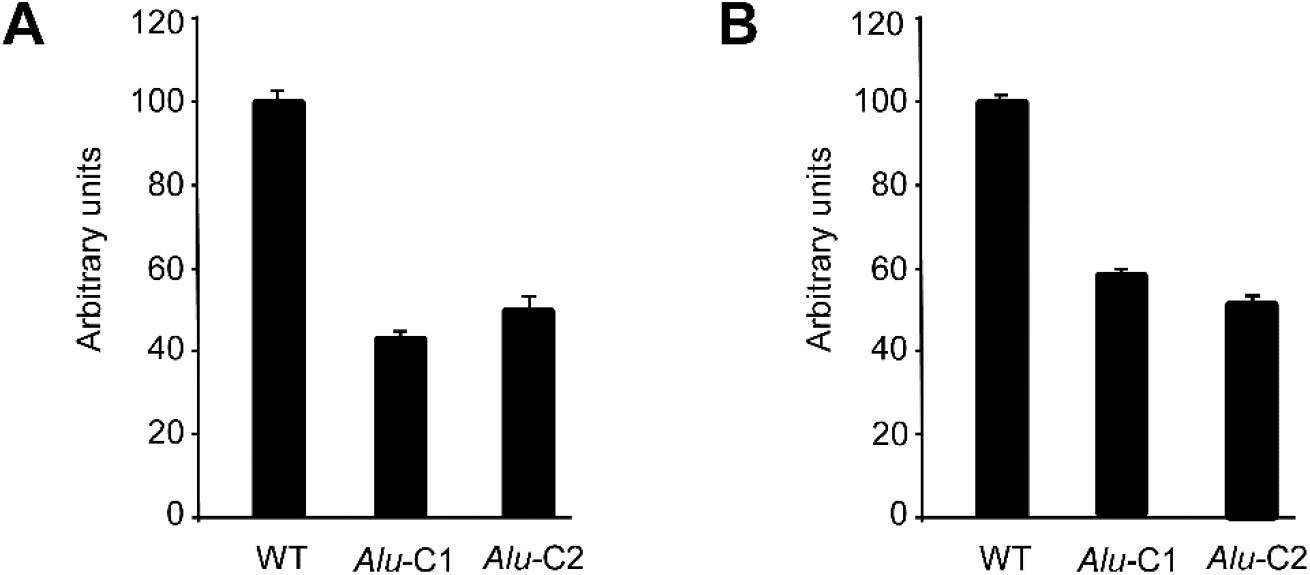
3’-UTR luciferase reporter assay outcomes. Transfections were performed in HEK293T (**A**) and COLO-357 (**B**) cells. “WT” refers to the wild-type *SPINK1* 3’-UTR luciferase reporter vector, whereas “*Alu*-C1” and “*Alu*-C2” denote two distinct clones of the mutated *SPINK1* 3’-UTR luciferase reporter vector. Data are shown as the average ± standard deviation from three separate experiments, each carried out in triplicate.

### 3.2. Formulation of a new hypothesis: impact of *Alu*_Ins on long double-stranded RNA (dsRNA) structure formation with pre-existing *Alu* elements in the *SPINK1* gene

A limitation of the 3’-UTR reporter assay lies in its inherent inability to assess the potential impact of the studied variant on splicing. Splicing, a complex process preceding mRNA 3’ end formation and other post-transcriptional events, is fundamental to gene expression [27-29]. The gene’s primary sequence, encoding regulatory elements and secondary structures, establishes the groundwork for mRNA splicing outcomes.

RNA secondary structures, dependent upon the primary sequence, significantly influence alternative splicing [30, 31]. Alterations in pre-mRNA secondary structure are linked to the pathological effects of numerous disease-associated variants [32, 33]. The abundance of *Alu* elements in the human genome, with their sequence similarities, makes inverted *Alu* repeats a major source for dsRNA structure formation. Notably, there are 228,607 inverted *Alu* pairs within 1 kb of each other in genes from the NCBI Reference Sequence (RefSeq) database [34]. Genome-wide analyses reveal that dsRNA structures frequently form between inverted *Alu* elements located within a distance of <800 bp apart, although they can also occur over distances of several thousand bp [35]. Recent research utilizing the J2 antibody has directly mapped dsRNA structures from inverted *Alu* elements in 3’-UTRs [36]. Additionally, artificial *Alu* sequences inserted in opposite orientations within an intron of a three-exon-minigene formed base pairings, thereby altering the splicing pattern of the downstream exon [37]. Lastly, the emergence of circular RNAs (circRNAs), a class of RNAs identified recently [38], is thought to result from back-splicing mechanisms [39]. Such mechanisms often depend on secondary structures in pre-mRNAs, especially those formed by inverted *Alu* elements [40-42].

Based on these findings, we hypothesize that the *SPINK1 Alu*_Ins variant could induce the formation of long dsRNA structures with pre-existing *Alu* elements in the *SPINK1* gene, potentially impacting splicing, one of the initial stages of mRNA formation.

### 3.3. Discovery of two oppositely oriented *Alu* elements in intron 3 of *SPINK1* gene relative to *Alu*_Ins

To establish the plausibility of our new hypothesis, two specific conditions would need to be met: Firstly, there must be pre-existing *Alu* elements within the *SPINK1* gene locus. Secondly, these pre-existing elements must be oriented inversely compared to the *Alu*_Ins variant. Using the RepeatMasker tool (http://www.repeatmasker.org/), we identified two such pre-existing *Alu* elements within intron 3 of the *SPINK1* gene, designated as *Alu*1 and *Alu*2, respectively. As illustrated in Supplementary Figure S1 and Figure 1, both *Alu*1 and *Alu*2 exhibit the opposite orientation to the *Alu*_Ins variant, thereby supporting our hypothesis. Furthermore, we observed that the pre-existing *Alu*2 element and the pathogenic *Alu*_Ins variant are separated by <800 bp. This close proximity is noteworthy, as inverted *Alu* repeats within such short distances often lead to the formation of dsRNA structures [35], which aligns with our investigative premise.

### 3.4. Prediction of long dsRNA stem formation between *Alu*_Ins and *Alu*2 using RNAfold

We employed RNAfold (http://rna.tbi.univie.ac.at/) [22], with default settings, to assess the potential secondary structures of the pre-mRNA of the Mut (*Alu*_Ins) allele, with the WT allele used for comparison. Our analysis encompassed sequences from the beginning of exon 3 to the end of the 3’-UTR (for the exact sequences, see Supplementary Figures S3 and S4). The minimum free energy (MFE) structures obtained, depicted in Figure 3, reveal the presence of an extensive dsRNA stem, covering 259 bp in *Alu*2 and 270 bp in *Alu*_Ins (refer to Supplementary Figure S4 for further details).

**Figure 3.**
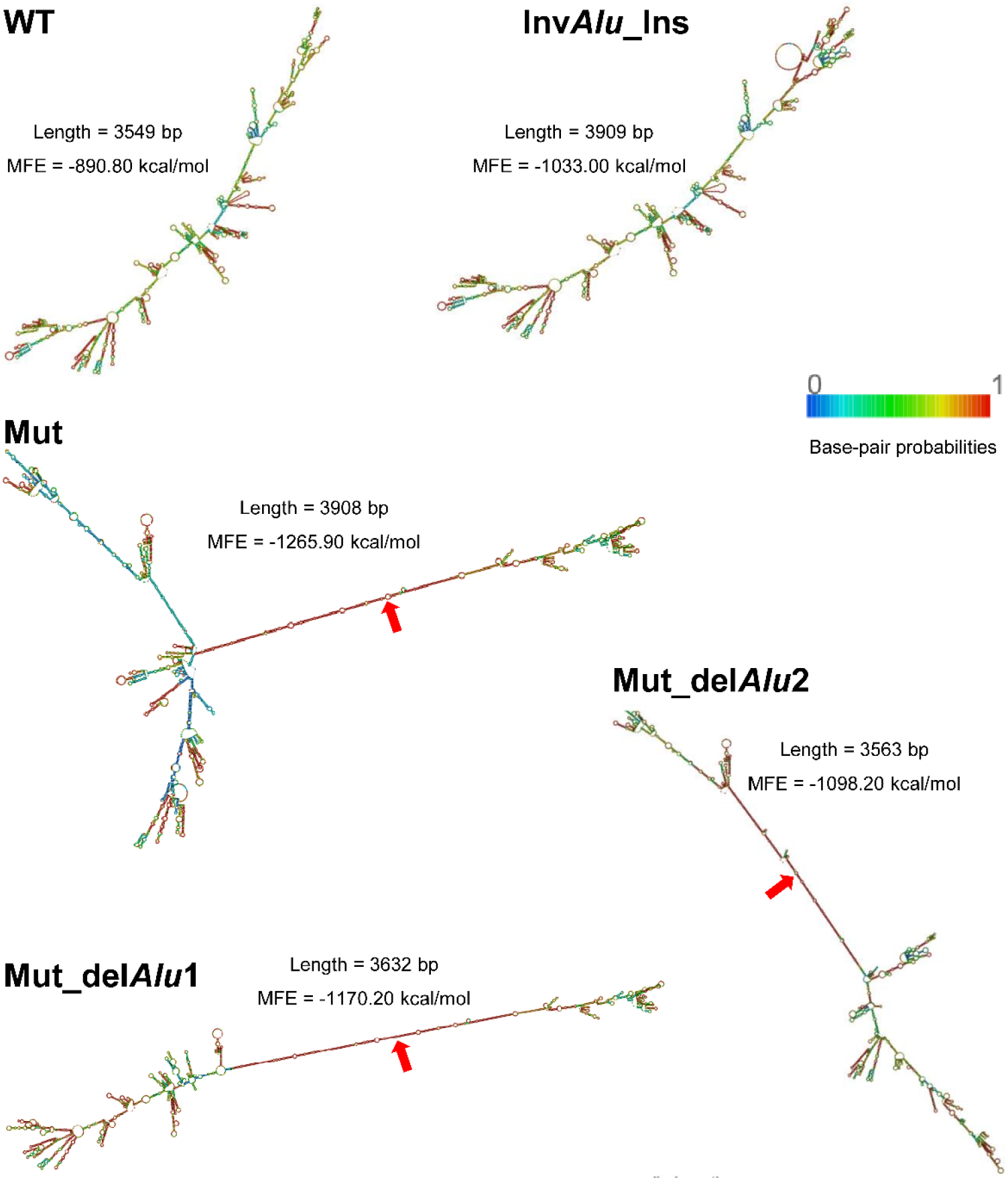
Minimum free energy (MFE) RNA secondary structures predicted by RNAfold for various *SPINK1* alleles. Refer to Figure 1 for a schematic representation of the different alleles. For each allele, the sequence used for prediction spans from the start of exon 3 to the end of the 3’-UTR of the *SPINK1* gene, with the corresponding length and MFE value specified within the Figure. Red arrows indicate the extended double-stranded RNA (dsRNA) stems formed between inverted *Alu* elements. The exact sequences employed in predicting RNA secondary structures are provided in Supplementary Figures S3-S7.

### 3.5. Restoration of *SPINK1* mRNA expression by disrupting long dsRNA stem formation

Given the established role of (i) inverted *Alu* repeats in forming long dsRNA stems and (ii) RNA secondary structure on gene splicing and expression, our findings would draw a direct link between the intricate dsRNA structure formed through the interaction of *Alu*2 and *Alu*_Ins and the absence of *SPINK1* expression. To delve deeper, we developed five new vectors for the FLGEA assay, notably introducing an inverted version of *Alu*_Ins, termed Inv*Alu*_Ins (see Figure 1). This modification aimed to prevent the formation of detrimental dsRNA structures by aligning all three *Alu* elements in the same direction, potentially restoring gene expression.

RNAfold analysis confirmed that Inv*Alu*_Ins does not form extensive dsRNA stems (Figure 3). Despite the nearly identical lengths of the Mut and Inv*Alu*_Ins sequences (3908 bp versus 3909 bp, respectively), noticeable differences were observed in their predicted MFE values: Mut exhibited -1265.90 kcal/mol, whereas Inv*Alu*_Ins showed -1033.00 kcal/mol. This discrepancy apparently arises from the extensive dsRNA stem within Mut. Furthermore, the significant difference in MFE between Mut and WT (−1265.90 kcal/mol vs. -890.80 kcal/mol) underscores the cumulative effects of increased sequence length and dsRNA structure in Mut. Generally, a more negative MFE indicates a stronger and more stable structure, whereas a less negative value suggests a weaker, less compact structure.

The Inv*Alu*_Ins vector, along with the other four newly designed vectors, was employed in the FLGEA assay together with the WT and Mut vectors. These assays were conducted under cycloheximide treatment, with the aim of revealing transcripts potentially masked by nonsense-mediated mRNA decay (NMD) in our previous research [8]. To begin, we shall discuss the results from the Inv*Alu*_Ins vector, comparing them to those from the WT and Mut vectors.

The reorientation of *Alu*_Ins within the Inv*Alu*_Ins vector effectively restored correctly spliced *SPINK1* transcripts. This was demonstrated by the detection of a distinct, clear band (identified as band G in Figure 4) in the RT-PCR analysis of HEK293T cells transfected with the Inv*Alu*_Ins vector; sequencing confirmed this band as representing correctly spliced transcripts containing the Ivs*Alu*_Ins sequence in the 3’-UTR. By contrast, cells transfected with the Mut vector displayed two distinct aberrant bands (labeled as bands A and B in Figure 4) and one faint band (referred to as band C in Figure 4). Sequencing identified the faint band C as correctly spliced transcripts that include *Alu*_Ins within the 3’-UTR.

**Figure 4.**
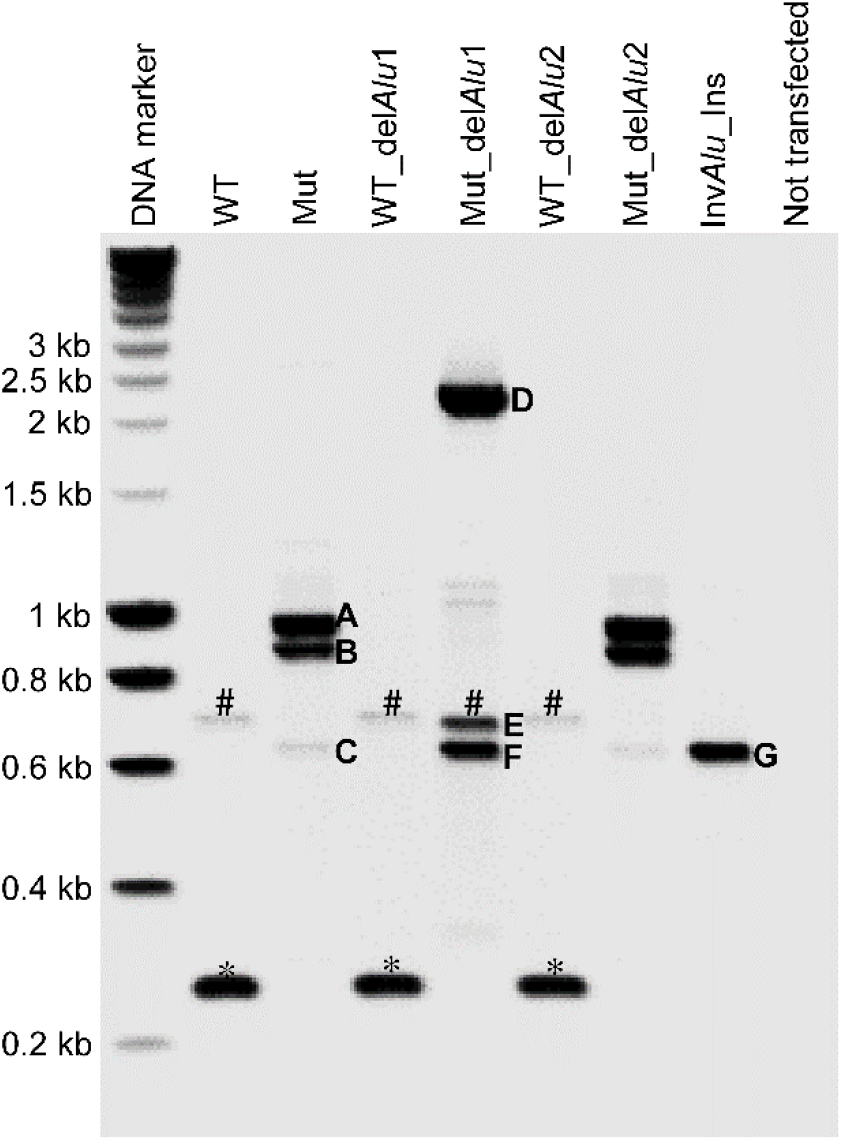
Outcomes of RT-PCR analysis on HEK293T cells transfected with various expression vectors. See Figure 1 for the schematic representation of the seven constructs. Bands marked with an asterisk indicate normally spliced wild-type *SPINK1* transcripts, while those with a hash represent non-specific amplifications. Bands C and F correspond to normally spliced transcripts containing *Alu*_Ins within the 3’-UTR, while band G corresponds to normally spliced transcripts that include Inv*Alu*_Ins within the 3’-UTR. Bands A, B, and D feature transcripts indicative of RNA template switching, facilitated by inverted *Alu* repeats. The characteristics of the three bands observed in Mut_del*Alu*2 closely mirror those in Mut.

The presence of the faintly spliced C band, alongside the two more pronounced aberrant bands (bands A and B), in cells transfected with the Mut vector in this study could be attributed to the effects of cycloheximide or differences in experimental conditions compared to our earlier study [8]. Nonetheless, the level of normally spliced transcripts in the Mut vector is notably low compared to either the co-amplified aberrant transcripts (bands A and B) observed in the Mut vector, the normally spliced transcripts detected with the WT vector (the leftmost band marked with an asterisk), or the normally spliced transcripts detected with the Inv*Alu*_Ins vector (band G). For example, the intensity of the faint C band from the Mut vector was measured to be only 4% of that of the G band from Inv*Alu*_Ins, as quantified using ImageJ.JS (https://ij.imjoy.io/). This substantially reduced level of normally spliced transcripts is consistent with the previously observed absence of *SPINK1* expression [8].

### 3.6. Further exploration of the impact of dsRNA formation on *SPINK1* expression by deletion of pre-existing *Alu*1 or *Alu*2 in the Mut context

The additional four vectors, designed with deletions of *Alu*1 or *Alu*2 elements within both the WT and Mut contexts, were utilized to further examine the effect of dsRNA formation on *SPINK1* expression. The results from the FLGEA assay, outlined in Figure 4, demonstrate that the WT context deletions, specifically WT_del*Alu*1 and WT_del*Alu*2, produced normally spliced WT transcripts, akin to those from the original WT vector, as evidenced by the second and third bands marked with asterisks. These observations imply that the deletions did not inadvertently create new splicing signals and confirm that neither *Alu*1 or *Alu*2 would have played a significant role in regulating *SPINK1* splicing within their original WT sequence context.

Using RNAfold, both the Mut_del*Alu*1 and Mut_del*Alu*2 alleles were predicted to develop significant dsRNA stem structures due to inverted *Alu* repeats, as illustrated in Figure 3. Specifically, interactions were noted between *Alu*2 and *Alu*_Ins in Mut_del*Alu*1 and between *Alu*1 and *Alu*_Ins in Mut_del*Alu*2 (see Supplementary Figures S6 and S7 for details). Consequently, despite being shorter than the Inv*Alu*_Ins sequence, which does not form extensive dsRNA structures, both Mut_del*Alu*1 and Mut_del*Alu*2 exhibited more negative MFE values compared to InvAlu_Ins (Figure 3).

Consistent with the predicted long dsRNA stem structures, we observed a marked decrease in normally spliced *SPINK1* transcripts with Mut_del*Alu*1 and almost total suppression with Mut_del*Alu*2 in the FLGEA assay. Specifically, after length adjustment, normally spliced *SPINK1* transcripts in Mut_del*Alu*1 (represented by band F in Figure 4) constituted about 90% of the level seen with the aberrantly spliced band D, signifying that under 45% of the total transcripts were normally spliced. By contrast, the RT-PCR results for Mut_del*Alu*2 closely matched those of the Mut allele. These results confirm the profound effect of dsRNA stem structures on gene splicing.

### 3.7. Aberrant RT-PCR bands harboring sequence features consistent with dsRNA formation

To identify the origins of the bands from the FLGEA, colony PCR analysis was conducted on the overall RT-PCR products for each vector, complemented by sequencing of excised bands when necessary. The results for the Mut vector—including the number of sequenced colonies and the characteristics of the RT-PCR products corresponding to bands A, B, and C—are shown in the left panel of Figure 5A. Sequences from over half (15 out of 29) of the informative Mut-derived colonies were consistent with normally spliced transcripts, associated with band C. The unusually high representation of band C, despite its faint appearance, is likely due to a cloning bias favoring shorter inserts (via ligation efficiency) during the cloning of RT-PCR products into the pcDNA3.1/V5-His TOPO vector.

**Figure 5.**
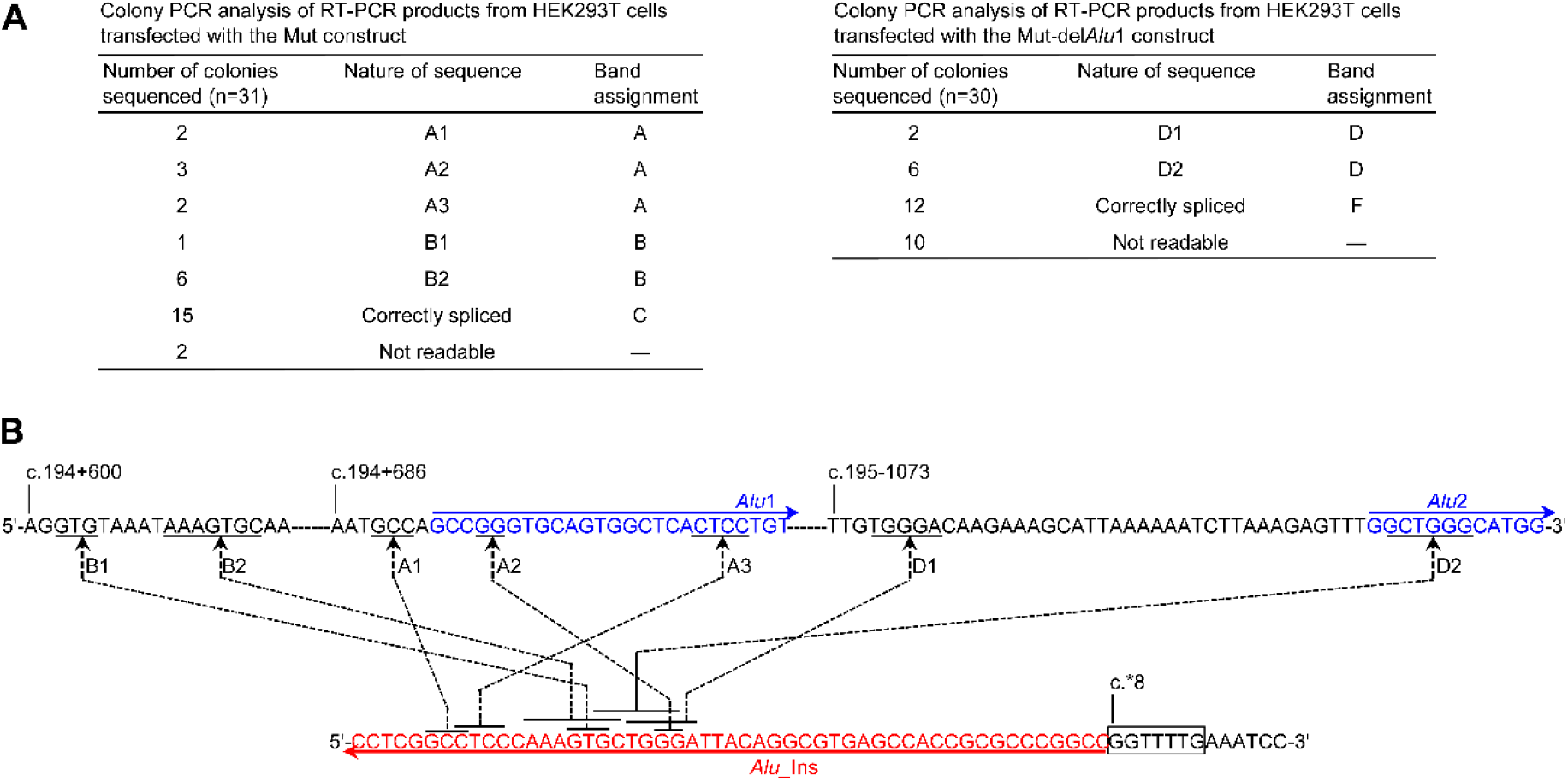
Results from colony PCR and sequencing analysis of FLGEA-derived RT-PCR bands. (**A**) Analysis of RT-PCR products from cells transfected with the Mut construct (left panel) or the Mut_del*Alu*1 construct (right panel). Refer to Figure 4 for the indicated bands. (**B**) Illustration of the presumed template switching events and their associated direct short repeats (indicated by underlines and overlines), responsible for the formation of aberrant transcripts (for more information, refer to the text). The boxed GGTTTG indicates the target site duplication. Nucleotide positions are designated according to the wild-type *SPINK1* sequence following the coding DNA sequence guidelines established by the Human Genome Variation Society [18]. *Abbreviations:* FLGEA, full-length gene expression assay; RT-PCR, reverse transcription-polymerase chain reaction.

From the remaining 14 informative Mut-derived colonies, five distinct products were identified. Three of these (A1, A2, and A3) corresponded to band A, and the other two (B1 and B2) to band B, as depicted in Figure 5A. The structure of these transcripts included exon 1, exon 2, exon 3, followed by a portion of the 5’ proximal part of intron 3, and a truncated 3’-UTR sequence. This pattern indicates that while introns 1 and 2 were correctly excised, intron 3 was partially retained, accompanied by significant internal deletions affecting both intron 3 and the 3’-UTR sequences. Importantly, the deletion junctions featured short direct repeats: one located within the *SPINK1 Alu*_Ins and another within the retained section of intron 3. These direct repeats within *SPINK1 Alu*_Ins were concentrated within a specific 20-bp segment, in contrast to those in intron 3, which were spread across a stretch less than 100-bp. This particular 20-bp segment within *SPINK1 Alu*_Ins is found near the 3’ end of *Alu*_Ins, whereas the less than 100-bp segment in intron 3 encompasses the 5’ end of *Alu*1, as illustrated in Figure 5B.

The identified sequence characteristics hint at a possible reverse transcriptase template switching mechanism [43], presumably instigated by dsRNA structures forming between *Alu*1 and *SPINK1 Alu*_Ins in the mutant pre-mRNA sequences. These structures might initially impede the proper splicing of intron 3 during pre-mRNA splicing and subsequently foster template switching during the reverse transcription of mutant mRNAs harboring dsRNA, thereby culminating in the emergence of bands A and B, as elaborated in the left panel of Figure 6. However, our attempts to verify the hypothesized full-length transcripts—those retaining intron 3 while accurately excluding introns 1 and 2—using different primer combinations were unsuccessful.

**Figure 6.**
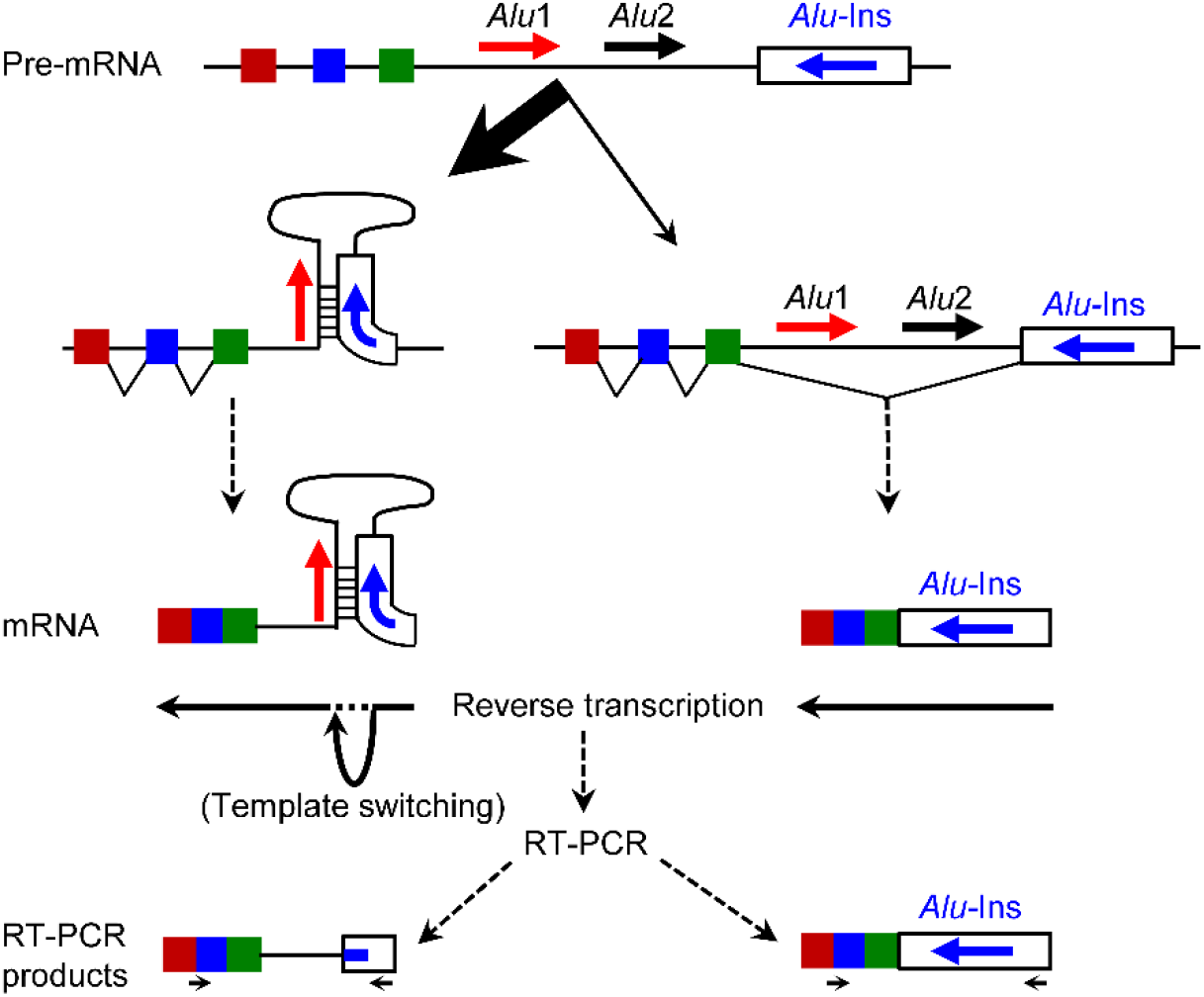
A schema to explain RT-PCR band formation in the FLGEA assay using the Mut vector. The components of the *SPINK1* genomic sequence are not to scale. The left panel demonstrates the production of the aberrant bands, while the right panel shows the creation of the faint band associated with normally spliced transcripts. In the bottom section, “RT-PCR products,” the facing arrows depict the approximate locations of the primers used for RT-PCR analysis: the forward primer spans the junction between exons 1 and 2, and the reverse primer corresponds to nucleotides c.29_46 of the *SPINK1* gene’s 3’-UTR. *Abbreviations:* FLGEA, full-length gene expression assay; RT-PCR, reverse transcription-polymerase chain reaction; 3’-UTR, 3’-untranslated region.

Colony PCR and sequencing investigations revealed that the three bands emerging from the deletion of *Alu*2 in the mutant *SPINK1* context (Mut_del*Alu*2) bore a close resemblance to those identified with the Mut vector (Figure 4). This congruence is logical, as the anomalous bands A and B in the Mut vector were likely spawned from the interplay between *Alu*1 and *Alu*_Ins. In the Mut_del*Alu*2 construct, *Alu*1 persists as the singular pre-existing *Alu* element poised for interaction with *Alu*_Ins.

For the Mut_del*Alu*1 vector, band F (Figure 4) typifies normally spliced transcripts. Efforts to sequence the gel-purified band E product were unsuccessful, yielding no discernible sequence after multiple attempts. Band D was found to comprise two disparate entities, D1 and D2, as depicted in the right panel of Figure 5A. It is noteworthy that D1 and D2 transcripts exhibit parallel attributes to the Mut-derived bands A and B. They correctly splice through introns 1 and 2, diverging at intron 3, which is followed by the 3’-UTR and suffers from a substantial internal deletion. The deletion junctions also feature short direct repeats: one within the *SPINK1 Alu*_Ins, located in the previously noted 20-bp segment, and the other within the sequence immediately upstream of the 5’ end of *Alu*2 (Figure 5B). This outcome is logical, given that in the Mut_del*Alu*1 construct, *Alu*2 is the sole remaining pre-existing *Alu* element that can interact with *Alu*_Ins. Thus, the mechanism postulated for the creation of Mut-derived bands (Figure 6) might similarly explain the formation of bands from Mut_del*Alu*1, emphasizing the significant influence of dsRNA stem structures on the splicing process.

## 4. Discussion

In this study, our objective was to determine the mechanisms underlying the complete loss of *SPINK1* gene expression associated with the homozygous *Alu* insertion (*Alu*_Ins) in the gene’s 3’-UTR [8]. Through a hypothesis-driven approach informed by current literature, we first identified pre-existing *Alu* elements within the third intron of the *SPINK1* gene, oriented in opposition to *Alu*_Ins. We then employed RNAfold predictions along with experimental data from a genomic context-preserving FLGEA assay, providing strong evidence that *Alu*_Ins’s harmful effects arise from its formation of extensive dsRNA structures with these pre-existing *Alu* sequences. Crucial evidence to the contrary came from the Inv*Alu*_Ins construct, an inverted version of the *Alu* insertion that aligns all three *Alu* elements in the same direction, thereby inhibiting dsRNA formation. This setup markedly contrasted with the Mut construct, which contains inverted *Alu* elements, sharply defining the role of *Alu* orientation and dsRNA structures in affecting gene expression.

Our findings are further reinforced by additional assays conducted with the FLGEA assay, focusing on the effects of deleting either the *Alu*1 or *Alu*2 elements. The outcomes of these deletion experiments were significant for two reasons. Firstly, the continued presence of inverted *Alu* elements in both Mut_del*Alu*1 and Mut_del*Alu*2 meant that a suppressive effect on gene expression was both expected and observed in the FLGEA assay. Secondly, and more intriguingly, the extent of suppression varied between the two. Contrary to initial expectations, Mut_del*Alu*1, but not Mut_del*Alu*2, showed a reduction in repressive impact on gene expression. This was surprising because *Alu*2 is physically closer to *Alu*_Ins than *Alu*1, and RNAfold predictions had indicated a potential interaction between *Alu*2 and *Alu*_Ins (Figure 3). This finding from the FLGEA assay implies that the *in vivo* effects of interactions between inverted *Alu* repeats on gene expression are complex, influenced by a variety of factors such as their proximity, sequence similarity, and possibly others. It is conceivable that, in a living organism, two *Alu* repeats positioned far apart and in opposite orientations could be brought into close proximity by complex RNA structures. It is also worth noting that in the Mut context, there were two pre-existing *Alu* elements, and RNAfold might have shown a preference for *Alu*2’s interaction with *Alu*_Ins over *Alu*1’s interaction. In summary, this comparative approach not only corroborates our initial findings but also deepens our understanding of the intricate molecular interactions involved. From this perspective, it is plausible to suggest that *in vivo*, the dsRNA structures involving the inverted *Alu* elements in the Mut allele might result from a network of interactions, potentially involving *Alu*1 and *Alu*_Ins, *Alu*2 and *Alu*_Ins, and even complex combinations of segments from *Alu*1, *Alu*2, and *Alu*_Ins.

The formation of extensive dsRNA structures between inverted *Alu* elements is anticipated to occur at the pre-mRNA level, hindering the normal splicing process of the *SPINK1* gene. Based upon current knowledge, this splicing issue should often result in the pre-mRNA, which contains dsRNA, not being processed into mature mRNA with a polyA tail, leading to its retention within the nucleus. Furthermore, even when dsRNA-containing mRNA is fully processed, it may still often be retained within the nucleus instead of being transported to the cytoplasm, as indicated by findings from a previous study [44]. With this context in mind, the aberrant RT-PCR bands observed in the FLGEA assays should be seen as being indicative of dsRNA-containing, fully processed, and aberrantly spliced mRNAs that have managed to be exported from the nucleus. Herein, we would like to emphasize that these aberrant RT-PCR bands represent genuine products under our experimental conditions, confirmed by the correct splicing out of introns 1 and 2 of the *SPINK1* gene. From the sequencing of these aberrant bands, we inferred that intron 3 of the *SPINK1* gene was retained in these mRNA sequences. However, within the constraints of our experimental setup, we were unable to capture the full-length transcripts that retained intron 3. Instead, what we detected were ‘template switching’ events that are characteristic of interactions involving the inverted *Alu* elements. Performing the reverse transcription at a higher temperature and using a higher quality reverse transcriptase could potentially enable the detection of full-length, aberrantly spliced transcripts.

Our study emphasizes the significance of utilizing appropriate model systems for functional analysis. Without the FLGEA assay, the detailed mechanisms behind the pathogenic effects of the *Alu* insertion (*Alu*_Ins) could not have been accurately identified. Indeed, informed by the current literature, one could logically infer a detrimental impact of *Alu*_Ins on the formation of the mRNA 3’ end and on various post-transcriptional events, as demonstrated by the 3’-UTR reporter assay.

## 5. Conclusions

While it has been well-documented that secondary structures can influence splicing and that inverted *Alu* elements are capable of forming such structures, our research marks the first instance of demonstrating an *Alu* insertion variant causing human genetic disease by creating extensive dsRNA structures with pre-existing *Alu* elements. Considering the widespread presence of *Alu* elements in the human genome and the potential for new *Alu* insertions at almost any locus, our findings carry significant implications for detecting and interpreting the impact of *Alu* element insertion events in genes associated with human disease. This suggests that the analysis of potential *Alu* insertions should extend beyond coding regions and proximal intronic sequences to include deep intronic and non-coding regions as well. Furthermore, the functional impact of *Alu* insertions needs to be assessed in terms of their distances from, and orientations relative to, existing *Alu* elements within the affected genes.

## Acknowledgments

We dedicate this work to the late Brice Felden, former Director of Inserm U1230 BRM (Bacterial RNAs and Medicine) at the Université de Rennes, Rennes, France. He provided expert insights into the formation of RNA secondary structures and their potential impact on gene expression regulation. Although sadly he was unable to see the project through to its completion, his influence was gratefully received and deeply felt.

## Author contributions

Conceptualization, E.M., C.F. and J.M.C.; study design, E.M. and J.M.C.; methodology, E.M., S.M., B.V. and J.M.C.; experiments, E.M., S.M., B.V.; data interpretation, E.M., S.M., B.V., D.N.C., C.F. and J.M.C.; writing— original draft preparation, J.M.C.; writing—review and editing, E.M., S.M., B.V., D.N.C., C.F. and J.M.C.; project administration, J.M.C.; funding acquisition, C.F. and J.M.C. All authors have read and approved the final version of the manuscript and agree to be accountable for all aspects of the work, ensuring that questions related to the accuracy or integrity of any part of the work are appropriately investigated and resolved.

## Data availability

All supporting data for this study are available within the article and its supplementary material.

## References

1. Lander ES, Linton LM, Birren B, Nusbaum C, Zody MC, Baldwin J, Devon K, Dewar K, Doyle M, FitzHugh W, et al: Initial sequencing and analysis of the human genome. Nature 2001, 409:860–921.

2. Kazazian HH, Jr., Moran JV: Mobile DNA in health and disease. N Engl J Med 2017, 377:361–370.

3. Feusier J, Watkins WS, Thomas J, Farrell A, Witherspoon DJ, Baird L, Ha H, Xing J, Jorde LB: Pedigree-based estimation of human mobile element retrotransposition rates. Genome Res 2019, 29:1567–1577.

4. Chen JM, Stenson PD, Cooper DN, Férec C: A systematic analysis of LINE-1 endonuclease-dependent retrotranspositional events causing human genetic disease. Hum Genet 2005, 117:411–427.

5. Hancks DC, Kazazian HH, Jr.: Roles for retrotransposon insertions in human disease. Mob DNA 2016, 7:9.

6. Chenais B: Transposable elements and human diseases: mechanisms and implication in the response to environmental pollutants. Int J Mol Sci 2022, 23.

7. Chen JM, Férec C, Cooper DN: LINE-1 endonuclease-dependent retrotranspositional events causing human genetic disease: mutation detection bias and multiple mechanisms of target gene disruption. J Biomed Biotechnol 2006, 2006:56182.

8. Venet T, Masson E, Talbotec C, Billiemaz K, Touraine R, Gay C, Destombe S, Cooper DN, Patural H, Chen JM, Férec C: Severe infantile isolated exocrine pancreatic insufficiency caused by the complete functional loss of the SPINK1 gene. Hum Mutat 2017, 38:1660–1665.

9. Pinto A, Cunha C, Chaves R, Butchbach MER, Adega F: Comprehensive in silico analysis of retrotransposon insertions within the survival motor neuron genes involved in spinal muscular atrophy. Biology (Basel) 2022, 11.

10. Chemparathy A, Guen YL, Zeng Y, Gorzynski J, Jensen T, Yang C, Kasireddy N, Talozzi L, Belloy ME, Stewart I, et al: A 3’UTR insertion is a candidate causal variant at the TMEM106B locus associated with increased risk for FTLD-TDP. medRxiv 2023.

11. Ohmuraya M, Hirota M, Araki M, Mizushima N, Matsui M, Mizumoto T, Haruna K, Kume S, Takeya M, Ogawa M, et al: Autophagic cell death of pancreatic acinar cells in serine protease inhibitor Kazal type 3-deficient mice. Gastroenterology 2005, 129:696–705.

12. Kume K, Masamune A, Kikuta K, Shimosegawa T: [-215G>A; IVS3+2T>C] mutation in the SPINK1 gene causes exon 3 skipping and loss of the trypsin binding site. Gut 2006, 55:1214.

13. Zou WB, Boulling A, Masson E, Cooper DN, Liao Z, Li ZS, Férec C, Chen JM: Clarifying the clinical relevance of SPINK1 intronic variants in chronic pancreatitis. Gut 2016, 65:884–886.

14. Chen JM, Lin JH, Masson E, Liao Z, Férec C, Cooper DN, Hayden M: The experimentally obtained functional impact assessments of 5’ splice site GT>GC variants differ markedly from those predicted. Curr Genomics 2020, 21:56–66.

15. Wu H, Lin JH, Tang XY, Marenne G, Zou WB, Schutz S, Masson E, Génin E, Fichou Y, Le Gac G, et al: Combining full-length gene assay and SpliceAI to interpret the splicing impact of all possible SPINK1 coding variants. Hum Genomics 2024, 18:21.

16. Yamamoto T, Nakamura Y, Nishide J, Emi M, Ogawa M, Mori T, Matsubara K: Molecular cloning and nucleotide sequence of human pancreatic secretory trypsin inhibitor (PSTI) cDNA. Biochem Biophys Res Commun 1985, 132:605–612.

17. Horii A, Kobayashi T, Tomita N, Yamamoto T, Fukushige S, Murotsu T, Ogawa M, Mori T, Matsubara K: Primary structure of human pancreatic secretory trypsin inhibitor (PSTI) gene. Biochem Biophys Res Commun 1987, 149:635–641.

18. den Dunnen JT, Dalgleish R, Maglott DR, Hart RK, Greenblatt MS, McGowan-Jordan J, Roux AF, Smith T, Antonarakis SE, Taschner PE: HGVS recommendations for the description of sequence variants: 2016 update. Hum Mutat 2016, 37:564–569.

19. Boulling A, Le Gac G, Dujardin G, Chen JM, Férec C: The c.1275A>G putative chronic pancreatitis-associated synonymous polymorphism in the glycoprotein 2 (GP2) gene decreases exon 9 inclusion. Mol Genet Metab 2010, 99:319–324.

20. Boulling A, Witt H, Chandak GR, Masson E, Paliwal S, Bhaskar S, Reddy DN, Cooper DN, Chen JM, Férec C: Assessing the pathological relevance of SPINK1 promoter variants. Eur J Hum Genet 2011, 19:1066–1073.

21. Zou WB, Boulling A, Masamune A, Issarapu P, Masson E, Wu H, Sun XT, Hu LH, Zhou DZ, He L, et al: No association between CEL-HYB hybrid allele and chronic pancreatitis in Asian populations. Gastroenterology 2016, 150:1558–1560 e1555.

22. Gruber AR, Lorenz R, Bernhart SH, Neubock R, Hofacker IL: The Vienna RNA websuite. Nucleic Acids Res 2008, 36:W70–74.

23. Chen JM, Férec C, Cooper DN: A systematic analysis of disease-associated variants in the 3’ regulatory regions of human protein-coding genes I: general principles and overview. Hum Genet 2006, 120:1–21.

24. Steri M, Idda ML, Whalen MB, Orru V: Genetic variants in mRNA untranslated regions. Wiley Interdiscip Rev RNA 2018, 9:e1474.

25. Griesemer D, Xue JR, Reilly SK, Ulirsch JC, Kukreja K, Davis JR, Kanai M, Yang DK, Butts JC, Guney MH, et al: Genome-wide functional screen of 3’UTR variants uncovers causal variants for human disease and evolution. Cell 2021, 184:5247–5260 e5219.

26. Romo L, Findlay SD, Burge CB: Regulatory features aid interpretation of 3’UTR variants. Am J Hum Genet 2024, 111:350–363.

27. Drexler HL, Choquet K, Churchman LS: Splicing kinetics and coordination revealed by direct nascent RNA sequencing through nanopores. Mol Cell 2020, 77:985–998 e988.

28. Gehring NH, Roignant JY: Anything but ordinary - emerging splicing mechanisms in eukaryotic gene regulation. Trends Genet 2021, 37:355–372.

29. Rogalska ME, Vivori C, Valcarcel J: Regulation of pre-mRNA splicing: roles in physiology and disease, and therapeutic prospects. Nat Rev Genet 2023, 24:251–269.

30. Solnick D: Alternative splicing caused by RNA secondary structure. Cell 1985, 43:667–676.

31. Clouet d’Orval B, d’Aubenton Carafa Y, Sirand-Pugnet P, Gallego M, Brody E, Marie J: RNA secondary structure repression of a muscle-specific exon in HeLa cell nuclear extracts. Science 1991, 252:1823–1828.

32. Buratti E, Baralle FE: Influence of RNA secondary structure on the pre-mRNA splicing process. Mol Cell Biol 2004, 24:10505–10514.

33. Chen JM, Férec C, Cooper DN: A systematic analysis of disease-associated variants in the 3’ regulatory regions of human protein-coding genes II: the importance of mRNA secondary structure in assessing the functionality of 3’ UTR variants. Hum Genet 2006, 120:301–333.

34. Daniel C, Silberberg G, Behm M, Ohman M: Alu elements shape the primate transcriptome by cis-regulation of RNA editing. Genome Biol 2014, 15:R28.

35. Bazak L, Levanon EY, Eisenberg E: Genome-wide analysis of Alu editability. Nucleic Acids Res 2014, 42:6876–6884.

36. Ku J, Lee K, Ku D, Kim S, Lee J, Bang H, Kim N, Do H, Lee H, Lim C, et al: Alternative polyadenylation determines the functional landscape of inverted Alu repeats. Mol Cell 2024.

37. Lev-Maor G, Ram O, Kim E, Sela N, Goren A, Levanon EY, Ast G: Intronic Alus influence alternative splicing. PLoS Genet 2008, 4:e1000204.

38. Kristensen LS, Andersen MS, Stagsted LVW, Ebbesen KK, Hansen TB, Kjems J: The biogenesis, biology and characterization of circular RNAs. Nat Rev Genet 2019, 20:675–691.

39. Welden JR, Stamm S: Pre-mRNA structures forming circular RNAs. Biochim Biophys Acta Gene Regul Mech 2019:194410.

40. Jeck WR, Sorrentino JA, Wang K, Slevin MK, Burd CE, Liu J, Marzluff WF, Sharpless NE: Circular RNAs are abundant, conserved, and associated with ALU repeats. RNA 2013, 19:141–157.

41. Liang D, Wilusz JE: Short intronic repeat sequences facilitate circular RNA production. Genes Dev 2014, 28:2233–2247.

42. Zhang XO, Wang HB, Zhang Y, Lu X, Chen LL, Yang L: Complementary sequence-mediated exon circularization. Cell 2014, 159:134–147.

43. Cocquet J, Chong A, Zhang G, Veitia RA: Reverse transcriptase template switching and false alternative transcripts. Genomics 2006, 88:127–131.

44. Chen LL, DeCerbo JN, Carmichael GG: Alu element-mediated gene silencing. EMBO J 2008, 27:1694–1705.

