## Supplementary Figure for "*Alu* insertion-mediated dsRNA structure formation with pre-existing *Alu* elements as a novel disease-causing mechanism"

CONTENTS

**Supplementary Figure S1.** Illustration of the *SPINK1* genomic sequence pertinent to this study in the context of the wild-type sequence.

**Supplementary Figure S2.** Schematic illustration of the primer pairs used to amplify distinct fragments for constructing the five additional expression vectors.

**Supplementary Figure S3.** Sequence of the wild-type *SPINK1* allele for RNAfold Prediction.

**Supplementary Figure S4.** Sequence of the mutant (*Alu*_Ins) *SPINK1* allele for RNAfold prediction.

**Supplementary Figure S5.** Sequence of the inverted allele of the *SPINK1 Alu*_Ins variant (Inv*Alu*_Ins) for RNAfold prediction.

**Supplementary Figure S6.** Sequence of Mut_del*Alu*1 (deletion of *Alu*1 within the mutant sequence context) for RNAfold Prediction.

**Supplementary Figure S7.** Sequence of Mut_del*Alu*2 (deletion of *Alu*2 within the mutant sequence context) for RNAfold Prediction.

c.1
|
**ATGAAGGTAACAGGCATCTTTCTTCTCAGTGCCTTGGCCCTGTTGAGTCTATCTG**gtaagtgttgcatattttc **Exon 1**
aaatttaaataaaactgttttgacctgttgctttgtgaagcacattatcttctagacttttgatgtagtctagtc
ttcgagagatgttttggacctaatgagatgaaataaaatcaacaggtaagaattatttttaagaggaattttta
acctactataaggaaaacaattctactagtaagaaattcccagaaataaaatggttttcctctattgatgtggct
gaccctttgttgggatatggtagagaaaaaagggtaaaaatatttaactttaagtatacttctcattctgtaaa
tgtgtaaggccaaattacaaattttaaattctcatgggattgcataaaatagataaatggcttccagttattcag
tataaaaggcataatttgactgattaaaaatttatttttttattgtataggtttcagtgtacttgaatcatgtat
agtttgtcagatttatccctgaatagctaatatatgtgatataaagggaatttttattttcaattcctggtatt
tcttattagtatctggaaatacaaaaattgcattttgtgtatttacattacacattttgaatgttttgaaagaaa
atattctgcttgaatagatggtgcacatatacctccaactgcatcttgacaatgttgacagacaactctgaaccc
gcagagccaagatcagtttccttttatcactggaaatggttaacattttcaacttcacctctcatgtagtgaag
gatcctgtggagagccaaatcagcactttgctaagaagaggaagaatattgtttgttttatttgctgaatattta
tggcaacagtcagtgttccctttccccctcctcatgcatgttaattaatacaaatagatgcctgtgatgagcacc
tgctcttcagtttttagactcaactagattctaaaagccagtgggcccccaacaacatcaagttctgactatccc
gggggaaattctccaagctcatagatgcctcactaattttcattacatttagatacaaagaactacctctttgct
cagatcacctcaaagattgatgatattaaaattaattttcaggaatgtttccttatttgattgctatcatattag
ttccctttcccttccatgaacttatccacagttatcatttcctatatgttgtcagcaaaacctctgctattttct
gggggaatagagattagctattttgactacctgtaaattcaacaaatgtgagttagacccatagtccatatgatg
caattgttggaaatagctctttgggaactgatagtgagttggcaaaagaactctatttttttcaacaatacttaca
aaagggaatttaaatgtaattgaaaaagctaactttgcatcagaagacctggttactgcttatgataactgtttc
acagctagttataaaaactctccatactttcatctgtaaactgagataaaaattatacttcactaattgtgagat
taaagatctagtaactggttttgagaatgtttgtgagtattcaaaagataatttgctcagtgcctgattcatttc
cagcagtaagtgcataatttctgttttcaaggaaatagatcaacttcctaaaacatcaaatcgagatactttggt
cataatcaaagttttactcaaacttttttaaggttttgccaaggaaggggtacaggaaaatcagtcagatactttt
ggaagattagagaaatcagaaagggtggggaatgaaagagcctagtaaagaagtcacagtctgcaatgaaagcag
agaattctgatgaagaatagatctgacttctttcatttaggacccaacttaccatatctgatttatttctag**GTA** **Exon 2**
**ACACTGGAGCTGACTCCCTGGGAAGAGAG**gtaaagagatatttgtaatttcttatttctcagactggaacagttt
gatccaacaaaaatgcagccttgctgtcacctttcagtttagcctgaagttaagaggagtatgaatgctagggag
ggttaacgtgagaaatgcaaatggtagtagtagttccaacccacaggcgtatattgaaggacaatttcactgatg
tataagcaagttccatatagtcctacagtgcttgctgtgaaggggtttttatatagactagtgatagctgtagac
aagtatttggggtggaagccttctcaacttcactccaagtagcccttcctccttacagccagtttgaccatgttg
gcatgcagctctcccaatcaccaaggtgtggctctaatttataatgaatgacagaaaataataaactaggttctc
cttctagaaaaatattctaactctaacctttcatattttttttgtttaagaggtaatttcagtaaatttggagac
cagaggaagaagtgtccctgttctcagtacatttatatctagtcagagacagctgatagaccaaaagttcatgaa
tgaatgcaatattacatttgtgataagtgttatgaaaaagttctgctgccgttaggaacttagcctagttagaaa
aatcagggagggtgctccgaagtaactgaggggttattttgagccaaaaaaaaaaaaatgagtgttagtaaaacaa
aactggtggggagatgtagtagaataaagtacttaagagctaaagctcaggaactctgactaggtttgaattctg
aatttaggaattccttagctgggtggccttggctaattttattgctttgtgcctcattatcttcacctgaagaag
aatataatacttagttcacaggatagttaggaggtttaaacagaataatatataaaaagcactagaacaatgtct
aggacaatgacaatcatttgatgattataaactcttactggagtagaatgcaaagacttccaacacccgacaatg
cctagtgcctttaagtgcctaatagaagaccagtgtgtgtggactacagtgtacaaggaagaacgtgccccaaga
tgtggccaacctgagagatgccagcacccgcagtaattgcaaacttgaacattatctcacccatccatgtaaatc
tcacaacaacactatgagataggtgggaaatggtaatatcatcactgttttgcagatgaactgactgagtttcag
aagggccataggacttactaatgtcacacagcttagaaatagcagaggcatgacttaaaacaaggttttctgtct

**Supplementary Figure S1.** Illustration of the *SPINK1* genomic sequence pertinent to this study in the context of the wild-type sequence. For the full-length gene expression assay (FLGEA), the sequence inserted into the pcDNA3.1/V5-His-TOPO vector extends from c.1 to c.*(300). For the 3'-UTR luciferase reporter assay, the sequence inserted into the pGL3 control vector extends from c.*1 to c.*(306) of the *SPINK1* gene. Bold uppercase letters: coding sequences. Blue highlights: *Alu1* and *Alu2* elements identified by RepeatMasker (N.B. both elements are in the same orientation as the sense strand of the *SPINK* gene). Grey shading: the 3'-UTR region. Double underline: target site duplication area relative to the disease-causing *Alu* insertion (*Alu*_Ins), which is in reverse orientation to the sense strand of the *SPINK1* gene. Upward arrow: *Alu*_Ins insertion site. Positions c.1 and c.*(306) correspond to genomic coordinates g.147831577 and g.147824355 in the GRCh38/hg38 chromosome 5 assembly, respectively. The *SPINK1* gene resides on the reverse strand of chromosome 5. The coding DNA sequence numbering follows the recommendations of the Human Genome Variation Society (HGVS).

ccagatagtaggttatttctcttacaacacacagtatcattctcccaatcacagttattccccagagaaataaaa
ccatttcagagattttgctatgaactcaagaatggagaataatgggaaatgattctgtttaattccatttttag**G**
**CCAAATGTTACAATGAACTTAATGGATGCACCAAGATATATGACCCTGTCTGTGGGACTGATGGAAATACTTATC** **Exon 3**
**CCAATGAATGCGTGTTATGTTTTGAAAATCG**gtgagtacaaacttgagtttcttttaaactatatatttaagtt

c.194

agttatcttcaagtgtactgataatatgaatctcaccccgagaaaagcaaactatttacttttccaaaaacagt
tatctctttcttattctccctttatatatttagcattaaatattatttttagaagtcacttgtatgataaaag
cctatatttttacagcaaaatagtcgatagcttggattttataaagtacacacaaaatacctttaattaaaaa
aattattagtggttgcaatgattgtgtttttaaagccctcagttatgcaaatataaattgtatagattatataa
ccaaaaatatattttacttatggatatgaaagccagtttcagtaaagcttcatttcctcagtgaccataagtag
gtgatgtgatacagagaaatagattcaacacagatttcctatctcactcagcaactgaaaccttagcatgtctca
agcatcatgctaatattccctgtcttacttgtgcttatgactaagaaaacatcatgagcatgtataggatggctt
cttataacttggggcaaatgccagaagatgaggtgtaaataaagtgcaacaggagtgaagcagagatacatattt
tttattaaaggggccaagaaaggatccatgaaaagaatagaatgccagccgggtgcagtggctcactcctgtaa
tcccagcactttgggaggccgatgcaggtggatcacctgagaccagcctggccaacatggtgaaaccccatggtg *Alu*1
gcgggcgcctataataccagctactcaggaggctgaggcagaagaatcgcttcaatctgggaggcagaggttgca
gtgagctgagattgtgccactgcactccaacctgggtgacaagagtgagactccatcttaaaaaaaaggaaaaaa
aaagaatagaatgccatttaaatagagttctgaattttggcaatatcaaaagagcatgccaggttaaataaaggt
gtggaatgaacaaagcaaataaaagggattccacgagtgactttagtttttctgagattgacttgatattaaaga
ctttgactcaggctcaaatgactgtgtaaatttccattatattctactgcataaattccctttctgtaatgcaa
accttcttcttgggctgttttaaaccattttggaaaattctcagagtttagcaagtattgtggaaataggacctc
tggatgagaggtctaatatcctgaagttttagacccaaaacattaggtattctactttgtaataaatctgtttcc
ataattaagacttgtaaaaagaatttgaagccacctgttagttttctcatgacccatgttcttgacattaagctc
aaacatcatatttccatccaatgggaggagaaagcaagagaacaacagctttgagacttgggaaatttccaagtg
ccgctcaacatttacagtgaaacattagaaaaatattagtagagggctggaacaccctatctgcttagcctctga
aacctccgagtttctgcttattctttctatttgttcaaaactatgtaaatatgggaggcacatttgtatgctaga
aaacatgacctctcttcctatatttctcagataagaaaatttaggtctagagacaaaatgttgctttctcaaata
caacatatttcctaatgtatgacattgcctttttcactgttgattaaaagtcccaaacttattacctagtaacta
aagccagacgctgaccctagtttgagacaagtagataataccaaacaagggcttgcaaatcagtctttttggtga
ttatgtgcttggcaaatatattcaataaacacttttctattccttctacgtcttcaaagggaatctgtaccggc
aaattcttttccataaacctgcatctccaatagaaccttctataatgatggaaatgttctatgtttatgctgtct
aatacatagccactagccacaaaacagtagccttaatgtggctaacactaacttaaatgtggctagtaaaactga
tcagctgaatatttaatattatctaaactaatttacatttacatagctatatgtggatagtgaatatatgaaact
agagtgaacagcatagcaactcttgtttattctgtttaggtgtgtgtgcttatgtatgctaactgatttggaaat
cagtattaatgagcagttaatgattgtgggacaagaaagcattaaaaaatcttaaagagtttggctgggcatggt
ggctcatgcctgtaatcccagcactttggaagtcccaggcaggtggatcacttttgttaggagtttgagaccagc
ctggccaacatggtaaaccccgtctctactaaaaatacaaatattagccaggcatggtggcattcacctgtaat *Alu*2
cccagctacttgggaggctgaggcaggagaatcacttgaacccgggaggtggaggctgcagtgagccaaggtcac
accactgcactccagcctgggcaacagagtgagactctgtcttaaaaaaagaaaaaaaagaaaaaaagaaaagaa
aaaaaaaagaaaaaaaagtatatggaagtcagatgtccttcagtaatagagaagtagaagaattttgtaagccc
gtacattcaaagaaaaaatggttagaaaaattaaagacctggaaacagtcttctagggaacaattataaaattta
gagatatttagtttagggaagagaaaactagtttggtaatggtagtgtggggcgaacagagaggttttatctgga
ggggaggatgtttagaactggcaggaatcatgtggaataaatatgtgatttacttttgtgagacctcacagatac
gaatccccagagctctaagatgaaaggtcaatcaattagaaggtccaacaatagctcaggcatcagctgagatgg
aaacctggtggcgaaggcagaggcatcaggagcaaaagatgcaacaatagttttgagtttaatttcaggggttca
ctgtaacaccccacgagatctgtaaaaactgtaaggacagatatttttcctttctgttgtatccttagtgtattc
aacagtaatcaataaatatttgatgaattaatgactaagtgaatgtataaatgaatgaaagacagagaaaagatt
ataaatctcaaacctctccaactttaaatgaagctgttatttttccccctgtttttctcccatagtcactttttc
atcagtgaagtttaagctgatatatttttttaatctctactgcag**GAAACGCCAGACTTCTATCCTCATTCAAA** **Exon 4**
**AATCTGGGCCTTGCTGA**gaaccaaggttttgaaatcccatcaggtcaccgcgaggcctgactggccttattgttg

c.240 c.*1

aataaatgtatctgaatatcccctgttgtttccatttgcttttcctcaaaggtatgtttgattataccagggtc

c.*81 c.*(82)

aggggactttggtaagactgagaatagaccaagttttaaaaacatcaggaatcctgtaatttcagacagggtttc
cctcataattcctttctttcctcatctttctcacatcctgaggtcactgccttagaaaatgtgattgcagatat
taaatggcagagcacagatgatc

c.*(300) c.*(306)

***Supplementary Figure S1*** *(continued)*

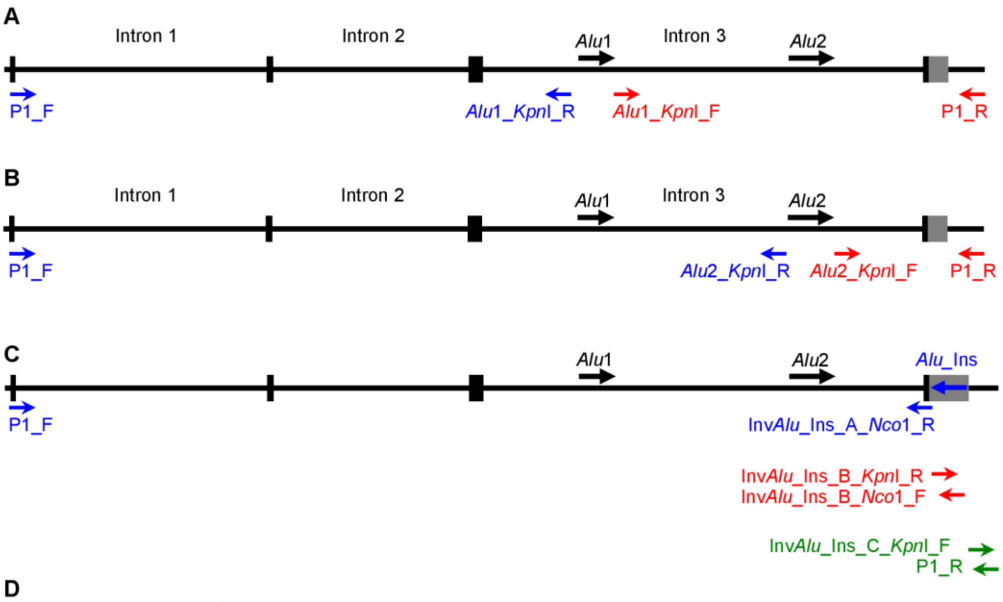


**D**

| Primer name | Primer sequence |
|---|---|
| P1_F | 5'-ATGAAGGTAACAGGCATCTTTCTTCTCAG-3' |
| *Alu*1_*KpnI*_R | 5'-CAGGAGTGAGCCACTGCACCCGGCTGGTACCATTCTATTCTTTTCATGGATCCT-3' |
| *Alu*1_*KpnI*_F | 5'- GAATAGAATGCCATTTAAATAGGTACCGTTCTGAATTTTGGCAATATCA-3' |
| *Alu*2_*KpnI*_R | 5'-CAGGCATGAGCCACCATGCCCAGGTACCAAACTCTTTAAGAT-3' |
| *Alu*2_*KpnI*_F | 5'-GTATATGGAAGTCAGATGGTACCTTCAGTAATAGAGAAG-3' |
| Inv*Alu*_Ins_A_*Nco*1_R | 5'- CGCCTGCCTAGCACCTCCATGGTTCTCAGCAAGGCCCAGATTTTTGAATGAGGATAGAAGTC-3' |
| Inv*Alu*_Ins_B_*KpnI*_R | 5'-CTTCTATCCTCATTCAAAAATCTGGGTACCTTGCTGAGAACCAAGGTTTTG-3' |
| Inv*Alu*_Ins_B_*Nco*1_F | 5'-CGGTGACCTGATGGGATTTCAAAACCATGGCCGGGCGCGGTGGCTCACGC-3' |
| Inv*Alu*_Ins_C_*KpnI*_F | 5'-CGGTTTTGAAATCCCATCAGGTACCACCGCGAGGCCTGACTGGC-3' |
| P1_R | 5'-CTGTGCTCTGCCATTTAATATCTGCAATC-3' |

**Supplementary Figure S2.** Schematic illustration of the primer pairs used to amplify distinct fragments for constructing the five additional expression vectors. (**A**) Two primer pairs (one indicated in blue, the other in red) were used to generate two fragments, whose ligation led to the deletion of *Alu*1 from the *SPINK1* genomic sequence. (**B**) Two primer pairs (one indicated in blue, the other in red) were used to generate two fragments, whose ligation resulted in the deletion of *Alu*2 from the *SPINK1* genomic sequence. In (**A**) and (**B**), deletions were performed in both WT and Mut sequence contexts (refer to Figure 1), with the former shown for illustrative purposes. (**C**) Within the context of the Mut (*Alu*_Ins) expression vector, three primer pairs (one indicated in blue, the second in red, the third in green) were used to generate three fragments, whose ligation resulted in the inversion of *Alu*-Ins (Inv*Alu*_Ins). (**D**) The sequences of the primers are provided, with the underlined portions denoting corresponding restriction enzyme recognition sites. The 5'-most ends of primers P1_F and P1_R correspond to nucleotide positions c.1 and c.*(300) of the *SPINK1* gene, respectively. In the primer names, 'F' denotes forward and 'R' signifies reverse.

**GCCAAAUGUUACAAUGAACUUAAUGGAUGCACCAAGAUAUAUGACCCUGUCUGUGGGACUGAUGGAAAUACUUAU** **Exon 3**
**CCCAAUGAAUGCGUGUUAUGUUUUGAAAAUCG**gugaguacaaacuugaguuucuuuuaaacuauauauuuuaagu
uaguuaucuucaaguguacugauaauaugaaucucaccccgagaaaagcaaacuauuuacuuuuuccaaaaacag
uuaucucuuucuuauucucccuuuuauauauuuagcauuaaauauuauuuuuuagaagucacuuguaugauaaaa
gccuauauuuuuuacagcaaaauagucgauagcuuggauuuuuauaaaguacacacaaaauaccuuuaauuaaaa
aaauuauuagugguugcaaugauuguguuuuuuaaagcccucaguuaugcaaauauaaauuguauagauuauaua
accaaaaauauauuuuuacuuauggauaugaaagccaguuucaguaaagcuucauuuccucagugaccauaagua
ggugaugugauacagagaaauagauucaacacagauuuccuaucucacucagcaacugaaaccuuagcaugucuc
aagcaucaugcuaauauucccugucuuacuugugcuuaugacuaagaaaacaucaugagcauguauaggauggcu
ucuuauaacuugggcaaaugccagaagaugagguguaaauaaagugcaacaggagugaagcagagauacauauu
uuuuauuaaaaggggccaagaaaggauccaugaaaagaauagaaugccagccgggugcaguggcucacuccugua
aucccagcacuuugggaggccgaugcagguggaucaccugagaccagccuggccaacauggugaaaccccauggu *Alu*1
ggcgggcgccuauaauaccagcuacucaggaggcugaggcagaagaaucgcuucaaucugggaggcagagguugc
agugagcugagauugugccacugcacuccaaccugggugacaagagugagacuccaucuuaaaaaaaaggaaaaa
aaaagaauagaaugccauuuaaauagaguucugaauuuggcaauaucaaaagagcaugccagguuaaauaaagg
uguggaaugaacaaagcaaauaaaagggauuccacgagugacuuuaguuuucugagauugacuugauauuaaag
acuuugacucaggcucaaaugacuguguaaauuuccauuauauucuacugcauaaauucccuuuucuguaaugca
aaccuucuucuugggcuguuuuaaaccauuuuggaaaauucucagaguuuagcaaguauuguggaaauaggaccu
cuggaugagaggucuaauauccugaaguuuuagacccaaaacauuagguauucuacuuuguaauaaaucuguuuc
cauaauuaagacuuguaaaaagaauuugaagccaccuguuaguuucucaugacccauguucuugacauuaagcu
caaacaucauauuuccauccaaugggaggagaaagcaagagaacaacagcuuugagacuugggaaauuuccaagu
gccgcucaacauuuacagugaaacauuagaaaaauauuaguagagggcuggaacacccuaucugcuuagccucug
aaaccuccgaguuucugcuuauucuuucuauuuguucaaaacuauguaaauaugggaggcacauuuguaugcuag
aaaacaugaccucucuuccuauauuucucagauaagaaaauuuaggucuagagacaaaauguugcuuucucaaau
acaacauauuuccuaauguaugacauugccuuuuucacuguugauuaaaagucccaaacuuauuaccuaguaacu
aaagccagacgcugacccuaguuugagacaaguagauaauaccaaacaagggcuugcaaaucagucuuuuuggug
auuaugugcuuggcaaauauauucaauaaacacuuuuucuauuccuucuacgucuucaaagggaaucuguaccgg
caaauucuuuuccauaaaccugcaucuccaauagaaccuucuauaaugauggaaauguucuauguuuaugcuguc
uaauacauagccacuagccacaaaacaguagccuuaauguggcuaacacuaacuuaaauguggcuaguaaaacug
aucagcugaauauuuaauauuaucuaaacuaauuuacauuuacauagcuauauguggauagugaauauaugaaac
uagagugaacagcauagcaacucuuguuuauucuguuuagguguguguguguuauguaugcuaacugauuuggaaa
ucaguauuaaugagcaguuaaugauugugggacaagaaagcauuaaaaaaucuuaaagaguuuggcugggcaugg
uggcucaugccuguaaucccagcacuuuggaagucccaggcagguggaucacuuuuguuaggaguuugagaccag
ccuggccaacaugguaaaaccccgucucuacuaaaaauacaaauauuagccaggcaugguggcauucaccuguaa *Alu*2
ucccagcuacuugggaggcugaggcaggagaaucacuugaacccgggagguggaggcugcagugagccaagguca
caccacugcacuccagccugggcaacagagugagacucugucuuaaaaaaagaaaaaaaagaaaaaaagaaaaga
aaaaaaaaagaaaaaaaaguauauggaagucagauguccuucaguaauagagaaguagaagaauuuuuguaagcc
cguacauucaaagaaaaaaugguuagaaaaauuaaagaccuggaaacagucuucuagggaacaauuauaaaauuu
agagauauuuaguuuagggaagagaaaacuaguuugguaaugguagugugggcgaacagagagguuuuaucugg
aggggaggauguuuagaacuggcaggaaucauguggaauaaauaugugauuuacuuuugugagaccucacagaua
cgaaucccagagcucuaagaugaaaggucaaucaauuagaagguccaacaauagcucaggcaucagcugagaug
gaaaccugguggcgaaggcagaggcaucaggagcaaaagaugcaacaauaguuugaguuuaauuucaggguuc
acuguaacaccccacgagaucuguaaaaacuguaaggacagauauuuuuccuuucuguuguauccuuaguguauu
caacaguaaucaauaaauauuugaugaauuaaugacuaagugaauguauaaaugaaugaaagacagagaaaagau
uauaaaucucaaaccucuccaacuuuaaaugaagcuguuauuuuccccuguuuucucccauagucacuuuuu
caucagugaaguuuaagcugauauauuuuuuuaaucucuacugcag**GAAACGCCAGACUUCUAUCCUCAUUCAA**
**AAAUCUGGGCCUUGCUGA**gaaccaagguuuugaaaucccaucaggucaccgcgaggccugacuggccuuauuguu **Exon 4**
gaauaaauguaucugaauaucccc

**Supplementary Figure S3.** Sequence of the wild-type *SPINK1* allele for RNAfold Prediction. This sequence spans from the start of exon 3 to the end of the 3'-untranslated region (3'-UTR). Coding sequences are in bold uppercase letters. Blue highlights indicate *Alu*1 and *Alu*2 elements as identified by RepeatMasker. The 3'-UTR region is marked with grey shading.

**GCCAAAUGUUACAAUGAACUUAAUGGAUGCACCAAGAUAUAUGACCCUGUCUGUGGGACUGAUGGAAAUACUUAU** **Exon 3**
**CCCAAUGAAUGCGUGUUAUGUUUUGAAAAUCG**gugaguacaaacuugaguuucuuuuaaacuauauauuuuaagu
uaguuaucuucaaguguacugauaauaugaaucucaccccgagaaaagcaaacuauuuacuuuuuccaaaaacag
uuaucucuuucuuauucucccuuuuauauauuuagcauuaaauauuauuuuuuagaagucacuuguaugauaaaa
gccuauauuuuuuacagcaaaauagucgauagcuuggauuuuuauaaaguacacacaaaauaccuuuaauuaaaa
aaauuauuagugguugcaaugauuguguuuuuuaaagcccucaguuaugcaaauauaaauuguauagauuauaua
accaaaaauauauuuuuacuuauggauaugaaagccaguuucaguaaagcuucauuuccucagugaccauaagua
ggugaugugauacagagaaauagauucaacacagauuuccuaucucacucagcaacugaaaccuuagcaugucuc
aagcaucaugcuaauauucccugucuuacuugugcuuaugacuaagaaaacaucaugagcauguauaggauggcu
ucuuauaacuuggggcaaaugccagaagaugagguguaaauaaagugcaacaggagugaagcagagauacauauu
uuuuauuaaaaggggccaagaaaggauccaugaaaagaauagaaugccagccgggugcaguggcucacuccugua
aucccagcacuuugggaggccgaugcagguggaucaccugagaccagccuggccaacauggugaaaccccauggu *Alu*1
ggcgggcgccuauaauaccagcuacucaggaggcugaggcagaagaaucgcuucaaucugggaggcagagguugc
agugagcugagauugugccacugcacuccaaccugggugacaagagugagacuccaucuuaaaaaaaaggaaaaa
aaaagaauagaaugccauuuaaauagaguucugaauuuggcaauaucaaaagagcaugccagguuaaauaaagg
uguggaaugaacaaagcaaauaaaagggauuccacgagugacuuuaguuuucugagauugacuugauauuaaag
acuuugacucaggcucaaaugacuguguaaauuuccauuauauucuacugcauaaauucccuuuucuguaaugca
aaccuucuucuugggcuguuuuaaaccauuuuggaaaauucucagaguuuagcaaguauuguggaaauaggaccu
cuggaugagaggucuaauauccugaaguuuuagacccaaaacauuagguauucuacuuuguaauaaaucuguuuc
cauaauuaagacuuguaaaaagaauuugaagccaccuguuaguuuucucaugacccauguucuugacauuaagcu
caaacaucauauuuccauccaaugggaggagaaagcaagagaacaacagcuuugagacuugggaaauuuccaagu
gccgcucaacauuuacagugaaacauuagaaaaauauuaguagagggcuggaacacccuaucugcuuagccucug
aaaccuccgaguuucugcuuauucuuucuauuuguucaaaacuauguaaauaugggaggcacauuuguaugcuag
aaaacaugaccucucuuccuauauuucucagauaagaaaauuuaggucuagagacaaaauguugcuuucucaaau
acaacauauuuccuaauguaugacauugccuuuuucacuguugauuaaaagucccaaacuuauuaccuaguaacu
aaagccagacgcugacccuaguuugagacaaguagauaauaccaaacaagggcuugcaaaucagucuuuuuggug
auuaugugcuuggcaaauauauucaauaaacacuuuuucuauuccuucuacgucuucaaagggaaucuguaccgg
caaauucuuuuccauaaaccugcaucuccaauagaaccuucuauaaugauggaaauguucuauguuuaugcuguc
uaauacauagccacuagccacaaaacaguagccuuaauguggcuaacacuaacuuaaauguggcuaguaaaacug
aucagcugaauauuuaauauuaucuaaacuaauuuacauuuacauagcuauauguggauagugaauauaugaaac
uagagugaacagcauagcaacucuuguuuauucuguuuagguguguguguggcuuauguaugcuaacugauuuggaaa
ucaguauuaaugagcaguuaaugauugugggacaagaaagcauuaaaaaaucuuaaagaguuuggcugggcaugg
uggcucaugccuguaaucccagcacuuuggaagucccaggcagguggaucacuuuuguuaggaguugagaccag
ccuggccaacaugguaaaaccccgucucuacuaaaaauacaaauauuagccaggcaugguggcauucaccuguaa *Alu*2
ucccagcuacuugggaggcugaggcaggagaaucacuugaacccgggagguggaggcugcagugagccaagguca
caccacugcacuccagccugggcaacagagugagacucugucuuaaaaaaagaaaaaaaagaaaaaaagaaaaga
aaaaaaaaagaaaaaaaaguauauggaagucagauguccuucaguaauagagaaguagaagaauuuuuguaagcc
cguacauucaaagaaaaaaugguuagaaaaauuaaagaccuggaaacagucuucuagggaacaauuauaaaauuu
agagauauuuaguuuagggaagagaaaacuaguuugguaaugguagugugggcgaacagagagguuuuaucugg
aggggaggauguuuagaacuggcaggaaucauguggaauaaauaugugauuuacuuuugugagaccucacagaua
cgaaucccccagagcucuaagaugaaaggucaaucaauuagaagguccaacaauagcucaggcaucagcugagaug
gaaaccugguggcgaaggcagaggcaucaggagcaaaagaugcaacaauaguuugaguuuaauuucaggguuc
acuguaacaccccacgagaucuguaaaaacuguaaggacagauauuuuuccuuucuguuguauccuuaguguauu
caacaguaaucaauaaauauuugaugaauuaaugacuaagugaauguauaaaugaaugaaagacagagaaaagau
uauaaaucucaaaccucuccaacuuuaaaugaagcuguuauuuuuccccuguuuuucucccauagucacuuuuu
caucagugaaguuuaagcugauauauuuuuuuuaaucucuacugcag**GAAACGCCAGACUUCUAUCCUCAUUCAA** **Exon 4**
**AAAUCUGGGCCUUGCUGA**gaaccaagguuuuguuauuauuauuauuuuuuuuuuuuuuuuuuuuuuuuuuuuaaa
aauuuuuuuuuuuuuuuuugagacggagucucgcucugucgcccaggccggacugcggacugcaguggcgcaauc
ucggcucacugcaagcuccgcuucccggguucacgccauucuccugccucagccuccccaguagcugggacuaca
ggcgcccgccaccgcgcccggcuaauuuuuuuguauuuuuaguagagacgggguuucaccuuguuagccaggau
ggucucaaucuccugaccucaugauccacccgccucggccucccaaagugcugggauuacaggcgugagccaccg
cgcccggccgguuuugaaaucccaucaggucaccgcgaggccugacuggccuuauuguugaauaaauguaucuga
auaucccc

**Supplementary Figure S4.** Sequence of the mutant (*Alu*_Ins) *SPINK1* allele for RNAfold prediction. This sequence covers from the beginning of exon 3 to the end of the 3'-untranslated region (3'-UTR). Coding sequences are presented in bold uppercase letters. Blue highlights denote the *Alu*1 and *Alu*2 elements, as characterized by RepeatMasker. The 3'-UTR region is distinguished by grey shading. Sequences flanking the *Alu* insertion (*Alu*_Ins) variant, which are highlighted in red, are singly underlined to indicate target site duplications. Sequences within the inverted *Alu*2 and *Alu*_Ins repeats, anticipated to form extensive stem structures as predicted by RNAfold, are doubly underlined.

**GCCAAAUGUUACAAUGAACUUAAUGGAUGCACCAAGAUAUAUGACCCUGUCUGUGGGACUGAUGGAAAUACUUAU** **Exon 3**
**CCCAAUGAAUGCGUGUUAUGUUUUGAAAAUCG**gugaguacaaacuugaguuucuuuuaaacuauauauuuuaagu
uaguuaucuucaaguguacugauaauaugaaucucaccccgagaaaagcaaacuauuuacuuuuuccaaaaacag
uuaucucuuucuuauucucccuuuuauauauuuagcauuaaauauuauuuuuuagaagucacuuguaugauaaaa
gccuauauuuuuuacagcaaaauagucgauagcuuggauuuuuauaaaguacacacaaaauaccuuuaauuaaaa
aaauuauuagugguugcaaugauuguguuuuuuaaagcccucaguuaugcaaauauaaauuguauagauuauaua
accaaaaauauauuuuuacuuauggauaugaaagccaguuucaguaaagcuucauuuccucagugaccauaagua
ggugaugugauacagagaaauagauucaacacagauuuccuaucucacucagcaacugaaaccuuagcaugucuc
aagcaucaugcuaauauucccugucuuacuugugcuuaugacuaagaaaacaucaugagcauguauaggauggcu
ucuuauaacuuggggcaaaugccagaagaugagguguaaauaaagugcaacaggagugaagcagagauacauauu
uuuuauuaaaaggggccaagaaaggauccaugaaaagaauagaaugccagccgggugcaguggcucacuccugua
aucccagcacuuugggaggccgaugcagguggaucaccugagaccagccuggccaacauggugaaaccccauggu *Alu*1
ggcgggcgccuauaauaccagcuacucaggaggcugaggcagaagaaucgcuucaaucugggaggcagagguugc
agugagcugagauugugccacugcacuccaaccugggugacaagagugagacuccaucuuaaaaaaaaggaaaaa
aaaagaauagaaugccauuuaaauagaguucugaauuuuggcaauaucaaaagagcaugccagguuaaauaaagg
uguggaaugaacaaagcaaauaaaaggauuccacgagugacuuuaguuuuucugagauugacuugauauuaaag
acuuugacucaggcucaaaugacuguguaaauuuccauuauauucuacugcauaaauucccuuuucuguaaugca
aaccuucuucuugggcuguuuuaaaccauuuuggaaaauucucagaguuuagcaaguauuguggaaauaggaccu
cuggaugagaggucuaauauccugaaguuuuagacccaaaacauuagguauucuacuuuguaauaaaucuguuuc
cauaauuaagacuuguaaaaagaauuugaagccaccuguuaguuuucucaugacccauguucuugacauuaagcu
caaacaucauauuuccauccaaugggaggagaaagcaagagaacaacagcuuugagacuugggaaauuuccaagu
gccgcucaacauuuacagugaaacauuagaaaaauauuaguagagggcuggaacacccuaucugcuuagccucug
aaaccuccgaguuucugcuuauucuuucuauuuguucaaaacuauguaaauaugggaggcacauuuguaugcuag
aaaacaugaccucucuuccuauauuucucagauaagaaaauuuaggucuagagacaaaauguugcuuucucaaau
acaacauauuuccuaauguaugacauugccuuuuucacuguugauuaaaagucccaaacuuauuaccuaguaacu
aaagccagacgcugacccuaguuugagacaaguagauaauaccaaacaagggcuugcaaaucagucuuuuuggug
auuaugugcuuggcaaauauauucaauaaacacuuuuucuauuccuucuacgucuucaaagggaaucuguaccgg
caaauucuuuuccauaaaccugcaucuccaauagaaccuucuauaaugauggaaauguucuauguuuaugcuguc
uaauacauagccacuagccacaaaacaguagccuuaauguggcuaacacuaacuuaaauguggcuaguaaaacug
aucagcugaauauuuaauauuaucuaaacuaauuuacauuuacauagcuauauguggauagugaauauaugaaac
uagagugaacagcauagcaacucuuguuuauucuguuuagguguguguguggcuuauguaugcuaacugauuuggaaa
ucaguauuaaugagcaguuaaugauugugggacaagaaagcauuaaaaaaucuuaaagaguuuggcugggcaugg
uggcucaugccuguaaucccagcacuuuggaagucccaggcagguggaucacuuuuguuaggaguuugagaccag
ccuggccaacaugguaaaaccccgucucuacuaaaaauacaaauauuagccaggcaugguggcauucaccuguaa *Alu*2
ucccagcuacuugggaggcugaggcaggagaaucacuugaacccgggagguggaggcugcagugagccaagguca
caccacugcacuccagccugggcaacagagugagacucugucuuaaaaaaagaaaaaaaagaaaaaaagaaaaga
aaaaaaaaagaaaaaaaaguauauggaagucagauguccuucaguaauagagaaguagaagaauuuuuguaagcc
cguacauucaaagaaaaaaugguuagaaaaauuaaagaccuggaaacagucuucuagggaacaauuauaaaauuu
agagauauuuaguuuagggaagagaaaacuaguuugguaaugguagugugggggcgaacagagagguuuuaucugg
aggggaggauguuuagaacuggcaggaaucauguggaauaaauaugugauuuacuuuugugagaccucacagaua
cgaaucccagagcucuaagaugaaaggucaaucaauuagaagguccaacaauagcucaggcaucagcugagaug
gaaaccugguggcgaaggcagaggcaucaggagcaaaagaugcaacaauaguuuugaguuuaauuucagggguuc
acuguaacaccccacgagaucuguaaaaacuguaaggacagauauuuuuccuuucuguuguauccuuaguguauu
caacaguaaucaauaaauauuugaugaauuaaugacuaagugaauguauaaaugaaugaaagacagagaaaagau
uauaaaucucaaaccucuccaacuuuaaaugaagcuguuauuuuuccccuguuuuucucccauagucacuuuuu
caucagugaaguuuaagcugauauauuuuuuuuaaucucuacugcag**GAAACGCCAGACUUCUAUCCUCAUUCAA**
**AAAUCUGGGCCUUGCUGA**gaaccaccauggccgggcgcgguggcucacgccuguaaucccagcacuuugggaggc **Exon 4**
cgaggcggguggaucaugaggucaggagauugagaccauccuggcuaacaaggugaaaccccgucucuacuaaaa
auacaaaaaaaauuagccgggcgcgguggcgggcgccuguagucccagcuacugggaggcugaggcaggagaau
ggcgugaacccgggaagcggagcuugcagugagccgagauugcgccacugcaguccgcaguccggccugggcgac
agagcgagacuccgucucaaaaaaaaaaaaaaaauuuuaaaaaaaaaaaaaaaaaaaaaaaaaaaauaauaau
aauaacaaaaccuugguucucagcaagguaccaccgcgaggccugacuggccuuauuguugaauaaauguaucug
aauaucccc

**Supplementary Figure S5.** Sequence of the inverted allele of the *SPINK1 Alu*_Ins variant (Inv*Alu*_Ins) for RNAfold prediction. This sequence extends from the start of exon 3 to the end of the 3'-untranslated region (3'-UTR). Coding sequences are depicted in bold uppercase letters. Blue highlights identify the *Alu*1 and *Alu*2 elements, as determined by RepeatMasker. The 3'-UTR region is marked with grey shading. Sequences highlighted in red represent the inverted version of the *Alu*_Ins variant.

**GCCAAAUGUUACAAUGAACUUAAUGGAUGCACCAAGAUAUAUGACCCUGUCUGUGGGACUGAUGGAAAUACUUAU** **Exon 3**
**CCCAAUGAAUGCGUGUUAUGUUUUGAAAAUCG**gugaguacaaacuugaguuucuuuuaaacuauauauuuuaagu
uaguuaucuucaaguguacugauaauaugaaucucaccccgagaaaagcaaacuauuuacuuuuuccaaaaacag
uuaucucuuucuuauucucccuuuuauauauuuagcauuaaauauuauuuuuuagaagucacuuguaugauaaaa
gccuauauuuuuuacagcaaaauagucgauagcuuggauuuuuauaaaguacacacaaaauaccuuuaauuaaaa
aaauuauuagugguugcaaugauuguguuuuuuaaagcccucaguuaugcaaauauaaauuguauagauuauaua
accaaaaauauauuuuuacuuauggauaugaaagccaguuucaguaaagcuucauuuccucagugaccauaagua
ggugaugugauacagagaaauagauucaacacagauuuccuaucucacucagcaacugaaaccuuagcaugucuc
aagcaucaugcuaauauucccugucuuacuugugcuuaugacuaagaaaacaucaugagcauguauaggauggcu
ucuuauaacuuggggcaaaugccagaagaugagguguaaauaaagugcaacaggagugaagcagagauacauauu
uuuuauuaaaaggggccaagaaaggauccaugaaaagaauagaauguaccguucugaauuuuggcaauaucaaa ~~*Alu1*~~
agagcaugccagguuaaauaaaggugugggaaugaacaaagcaaauaaaagggauuccacgagugacuuuaguuuu
ucugagauugacuugauauuaaagacuuugacucaggcucaaaugacuguguaaauuuccauuauauucuacugc
auaaauucccuuuucuguaaugcaaaccuucuucuugggcuguuuuaaaccauuuuggaaaauucucagaguuua
gcaaguauuguggaaauaggaccucuggaugagaggucuaauauccugaaguuuuagacccaaaacauuagguau
ucuacuuuguaauaaaucuguuuccauaauuaagacuuguaaaaagaauuugaagccaccuguuaguuuucucau
gacccauguucuugacauuaagcucaaacaucauauuuccauccaaugggaggagaaagcaagagaacaacagcu
uugagacuugggaaauuuccaagugccgcucaacauuuacagugaaacauuagaaaaauauuaguagagggcugg
aacacccuaucugcuuagccucugaaaccuccgaguuucugcuuauucuuucuauuuguucaaaacuauguaaau
augggaggcacauuuguaugcuagaaaacaugaccucucuucccuauauuucucagauaagaaaauuuaggucuag
agacaaaauguugcuuucucaaauacaacauauuuccuaauguaugacauugccuuuuucacuguugauuaaaag
ucccaaacuuauuaccuaguaacuaaagccagacgcugacccuaguuugagacaaguagauaauaccaaacaagg
gcuugcaaaucagucuuuuuggugauuaugugcuuggcaaauauauucaauaaacacuuuuucuauuccuucuac
gucuucaaagggaaucuguaccggcaaauucuuuuccauaaaccugcaucuccaauagaaccuucuauaaugaug
gaaauguucuauguuuaugcugucuaauacauagccacuagccacaaaacaguagccuuaauguggcuaacacua
acuuaaauguggcuaguaaaacugaucagcugaauauuuaauauuaucuaaacuaauuuacauuuacauagcuau
auguggauagugaauauaugaaacuagagugaacagcauagcaacucuuguuuauucuguuuaggugugugugcu
uauguaugcuaacugauuuggaaaucaguauuaaugagcaguuaaugauugugggacaagaaagcauuaaaaaau
cuuaaagaguuuggcugggcaugguggcucaugccuguaaucccagcacuuuggaagucccaggcagguggauca
cuuuuguuaggaguuugagaccagccuggccaacaugguaaaaccccgucucuacuaaaaauacaaauauuagcc
aggcaugguggcauucaccuguaaucccagcuacuugggaggcugaggcaggagaaucacuugaacccgggaggu *Alu2*
ggaggcugcagugagccaaggucacaccacugcacuccagccugggcaacagagugagacucugucuuaaaaaaa
gaaaaaaaagaaaaaaagaaaagaaaaaaaaaagaaaaaaaaguauauggaagucagauguccuucaguaauaga
gaaguagaagaauuuuuguaagcccguacauucaaagaaaaaaugguuagaaaaauuaaagaccuggaaacaguc
uucuagggaacaauuauaaaauuuagagauauuuaguuuagggaagagaaaacuaguuugguaaugguagugugg
ggcgaacagagagguuuuaucuggaggggaggauguuuagaacuggcaggaaucauguggaauaaauaugugauu
uacuuuugugagaccucacagauacgaauccccagagcucuaagaugaaaggucaaucaauuagaagguccaaca
auagcucaggcaucagcugagauggaaaccugguggcgaaggcagaggcaucaggagcaaaagaugcaacaauag
uuuugaguuuaauuucagggguucacuguaacaccccacgagaucuguaaaaacuguaaggacagauauuuuucc
uuucuguuguauccuuaguguauucaacaguaaucaauaaauauuugaugaauuaaugacuaagugaauguauaa
augaaugaaagacagagaaaagauuauaaaucucaaaccucuccaacuuuaaaugaagcuguuauuuuuccccccu
guuuuucucccauagucacuuuuucaucagugaaguuuaagcugauauauuuuuuuuaaucucuacugcag**GAAA** **Exon 4**
**CGCCAGACUUCUAUCCUCAUUCAAAAAUCUGGGCCUUGCUGA**gaaccaagguuuuguuauuauuauuauuuuuuu
uuuuuuuuuuuuuuuuuuuuuuaaaauuuuuuuuuuuuuuuuuuugagacggagucucgcucugucgcccaggccgg
acugcggacugcaguggcgcaaucucggcucacugcaagcuccgcuucccggguucacgccauucuccugccuca
gccuccccaguagcugggacuacaggcgcccgccaccgcgcccggcuaauuuuuuuguauuuuuaguagagacg
ggguuucaccuuguuagccaggauggucucaaucuccugaccucaugauccacccgccucggccucccaaagugc
ugggauuacaggcgugagccaccgcgcccggccgguuuugaaaucccaucaggucaccgcgaggccugacuggcc
uuauuguugaauaaauguaucugaauaucccc

**Supplementary Figure S6.** Sequence of Mut_del*Alu*1 (deletion of *Alu*1 within the mutant sequence context) for RNAfold Prediction. This sequence spans from the beginning of exon 3 to the conclusion of the 3'-untranslated region (3'-UTR). The original *Alu*1 element has been excised, and the novel sequence introduced at the deletion junction is shaded in yellow. Coding sequences are presented in bold uppercase letters. The remaining pre-existing *Alu*2 element within intron 3 is highlighted in blue. The 3'-UTR region is distinguished by grey shading. Sequences flanking the *Alu* insertion (*Alu*_Ins) variant, marked in red, are singly underlined to denote target site duplications. Additionally, sequences predicted by RNAfold to form extensive stem structures within the inverted *Alu*2 and *Alu*_Ins repeats are doubly underlined.

**GCCAAAUGUUACAAUGAACUUAAUGGAUGCACCAAGAUAUAUGACCCUGUCUGUGGGACUGAUGGAAAUACUUAU** Exon 3
**CCCAAUGAAUGCGUGUUAUGUUUUGAAAAUCG**gugaguacaaacuugaguuucuuuuaaacuauauauuuuaagu
uaguuaucuucaaguguacugauaauaugaaucucaccccgagaaaagcaaacuauuuacuuuuuccaaaaacag
uuaucucuuucuuauucucccuuuuauauauuuagcauuaaauauuauuuuuuagaagucacuuguaugauaaaa
gccuauauuuuuuacagcaaaauagucgauagcuuggauuuuuauaaaguacacacaaaauaccuuuaauuaaaa
aaauuauuagugguugcaaugauuguguuuuuuaaagcccucaguuaugcaaauauaaauuguauagauuauaua
accaaaaauauauuuuuacuuauggauaugaaagccaguuucaguaaagcuucauuuccucagugaccauaagua
ggugaugugauacagagaaauagauucaacacagauuuccuaucucacucagcaacugaaaccuuagcaugucuc
aagcaucaugcuaauauucccugucuuacuugugcuuaugacuaagaaaacaucaugagcauguauaggauggcu
ucuuauaacuuggggcaaaugccagaagaugagguguaaauaaagugcaacaggagugaagcagagauacauauu
uuuuauuaaaaggggccaagaaaggauccaugaaaagaauagaaugccagccgggugcaguggcucacuccugua
aucccagcacuuugggaggccgaugcagguggaucaccugagaccagccuggccaacauggugaaaccccauggu *Alu*1
ggcgggcgccuauaauaccagcuacucaggaggcugaggcagaagaaucgcuucaaucugggaggcagagguugc
agugagcugagauugugccacugcacuccaaccugggugacaagagugagacuccaucuuaaaaaaaaggaaaaa
aaaagaauagaaugccauuuaaauagaguucugaauuuuggcaauaucaaaagagcaugccagguuaaauaaagg
uguggaaugaacaaagcaaauaaaagggauuccacgagugacuuuaguuuuucugagauugacuugauauuaaag
acuuugacucaggcucaaaugacuguguaaauuuccauuauauucuacugcauaaauucccuuuucuguaaugca
aaccuucuucuugggcuguuuuaaaccauuuuggaaaauucucagaguuuagcaaguauuguggaaauaggaccu
cuggaugagaggucuaauauccugaaguuuuagacccaaaacauuagguauucuacuuuguaauaaaucuguuuc
cauaauuaagacuuguaaaaagaauuugaagccaccuguuaguuucucaugacccauguucuugacauuaagcu
caaacaucauauuuccauccaaugggaggagaaagcaagagaacaacagcuuugagacuugggaaauuuccaagu
gccgcucaacauuuacagugaaacauuagaaaaauauuaguagagggcuggaacacccuaucugcuuagccucug
aaaccuccgaguuucugcuuauucuuucuauuuguucaaaacuauguaaauaugggaggcacauuuguaugcuag
aaaacaugaccucucuuccuauauuucucagauaagaaaauuuaggucuagagacaaaauguugcuuucucaaau
acaacauauuuccuaauguaugacauugccuuuuucacuguugauuaaaagucccaaacuuauuaccuaguaacu
aaagccagacgcugacccuaguuugagacaaguagauaauaccaaacaagggcuugcaaaucagucuuuuuggug
auuaugugcuuggcaaauauauucaauaaacacuuuuucuauuccuucuacgucuucaaagggaaucuguaccgg
caaauucuuuuccauaaaccugcaucuccaauagaaccuucuauaaugauggaaauguucuauguuuaugcuguc
uaauacauagccacuagccacaaaacaguagccuuaauguggcuaacacuaacuuaaauguggcuaguaaaacug
aucagcugaauauuuaauauuaucuaaacuaauuuacauuuacauagcuauauguggauagugaauauaugaaac
uagagugaacagcauagcaacucuuguuuauucuguuuagguguguguguuaugcuaacugauuuggaaa
ucaguauuaaugagcaguuaaugauugugggacaagaaagcauuaaaaaaucuuaaagaguuugguaccuucagu ~~*Alu*2~~
aauagagaaguagaagaauuuuuguaagcccguacauucaaagaaaaaaugguuagaaaaauuaaagaccuggaa
acagucuucuagggaacaauuauaaaauuuagagauauuuaguuuagggaagagaaaacuaguuugguaauggua
guguggggcgaacagagagguuuuaucuggaggggaggauguuuagaacuggcaggaaucauguggaauaaauau
gugauuuacuuuugugagaccucacagauacgaauccccagagcucuaagaugaaaggucaaucaauuagaaggu
ccaacaauagcucaggcaucagcugagauggaaaccugguggcgaaggcagaggcaucaggagcaaaagaugcaa
caauaguuuugaguuuaauuucagggguucacuguaacaccccacgagaucuguaaaaacuguaaggacagauau
uuuuccuuucuguuguauccuuaguguauucaacaguaaucaauaaauauuugaugaauuaaugacuaagugaau
guauaaaugaaugaaagacagagaaaagauuauaaaucucaaaccucuccaacuuuaaaugaagcuguuauuuuu
ccccuguuuuucucccauagucacuuuuucaucagugaaguuuaagcugauauauuuuuuuuaaucucuacugc
ag**GAAACGCCAGACUUCUAUCCUCAUUCAAAAAUCUGGGCCUUGCUGA**gaaccaagguuuuguuauuauuauuau Exon 4
uuuuuuuuuuuuuuuuuuuuuuuuuuuuuuuaaaaauuuuuuuuuuuuuuuuugagacggagucucgcucugucgccca
ggccggacugcggacugcaguggcgcaaucucggcucacugcaagcuccgcuucccggguucacgccauucuccu
gccucagccuccccaguagcugggacuacaggcgcccgccaccgcgcccggcuaauuuuuuuguauuuuuagua
gagacgggguuucaccuuguuagccaggauggucucaaucuccugaccucaugauccacccgccucggccuccca
aagugcugggauuacaggcgugagccaccgcgcccggccgguuuugaaaucccaucaggucaccgcgaggccuga
cuggccuuauuguugaauaaauguaucugaauauccccc

**Supplementary Figure S7.** Sequence of Mut_del*Alu*2 (deletion of *Alu*2 within the mutant sequence context) for RNAfold Prediction. This sequence spans from the beginning of exon 3 to the conclusion of the 3'-untranslated region (3'-UTR). The original *Alu*2 element has been excised, and the novel sequence introduced at the deletion junction is shaded in yellow. Coding sequences are presented in bold uppercase letters. The remaining pre-existing *Alu*1 element within intron 3 is highlighted in blue. The 3'-UTR region is distinguished by grey shading. Sequences flanking the *Alu* insertion (*Alu*_Ins) variant, marked in red, are singly underlined to denote target site duplications. Additionally, sequences predicted by RNAfold to form extensive stem structures within the inverted *Alu*1 and *Alu*_Ins repeats are doubly underlined.
